# RNA-Binding and Prion Domains: The Yin and Yang of Phase Separation

**DOI:** 10.1101/2020.01.14.904383

**Authors:** Nieves Lorenzo Gotor, Alexandros Armaos, Giulia Calloni, R. Martin Vabulas, Natalia Sanchez de Groot, Gian Gaetano Tartaglia

**Affiliations:** Centre for Genomic Regulation (CRG), The Barcelona Institute for Science and Technology, Dr. Aiguader 88, 08003 Barcelona, Spain and Universitat Pompeu Fabra (UPF), 08003 Barcelona, Spain; Buchmann Institute for Molecular Life Sciences, Goethe University Frankfurt, Frankfurt am Main, Germany; Institute of Biophysical Chemistry, Goethe University Frankfurt, Frankfurt am Main, Germany; Institucio Catalana de Recerca i Estudis Avançats (ICREA), 23 Passeig Lluis Companys, 08010 Barcelona, Spain; Department of Biology ‘Charles Darwin’, Sapienza University of Rome, P.le A. Moro 5, Rome 00185, Italy; Department of Neuroscience and Brain Technologies, Istituto Italiano di Tecnologia, Via Morego 30, 16163, Genoa, Italy

## Abstract

Biomolecular condensates are membrane-less organelles mainly composed of proteins and RNAs that organize intracellular spaces and regulate biochemical reactions. The ability of proteins and RNAs to phase separate is encoded in their sequences, yet it is still unknown which domains drive the process and what are their specific roles. Here, we systematically investigated the human and yeast proteomes to find regions promoting biomolecular condensation. Using advanced computational models to predict the phase separation propensity of proteins, we designed a set of experiments to investigate the contributions of Prion-Like Domains (PrLDs) and RNA-Binding Domains (RBDs). We found that while just one PrLD is sufficient to drive protein condensation, multiple RBDs are needed to modulate the dynamicity of the assemblies. In the case of stress granule protein Pub1 we show that the PrLD promotes sequestration of protein partners and the RBD confers liquid-like behaviour to the condensate. Our work sheds light on the fine interplay between RBDs and PrLD to regulate formation of membrane-less organelles, opening up the avenue for their manipulation.

## Introduction

A correct functioning of cells requires a strict spatio-temporal regulation of molecular processes ^1^. To ensure such regulation, cells contain different organelles: some of them, such as vacuoles and lysosomes, are separated by membranes while others, such as stress granules (SGs) and processing-bodies (PBs), form when their constituent components condense in large assemblies ^2, 3, 4, 5^. In the past decades, high interest has been drawn to study the intracellular condensates due to their key role in regulatory processes and association with several human diseases ^6^.

Membrane-less organelles are diverse and versatile ^5, 7^. Currently, we know more than 10 different types of intracellular condensates involved in functions ranging from storage to transcriptional regulation ^5, 8, 9^. These assemblies, usually composed of proteins and RNAs, have liquid-like behaviour, adopt spherical shape and undergo deformation and fusion events ^3, 4, 10^. Their formation has been described to require a liquid-liquid phase separation process, in which the energies of inter-molecular interactions are greater than the entropy of being mixed with the solvent ^2^. In this process, components assemble by multiple dynamic contacts (known as multivalent interactions ^11, 12, 13^) allowing local accumulation, exchange of elements with the surrounding environment and formation of areas with different density and composition ^14, 15^. The different composition facilitates the development of sequential biological reactions within one specific biological condensate^15, 16, 17^. For example, the nucleolus has three immiscible sub-compartments that coordinate the sequential assembly of ribosomes ^15^.

Even within the same biomolecular condensate the material state and composition vary with the environmental conditions ^9^. Thus, a condensate change the dynamicity (solid- or liquid-like behaviour), depending on the type of contacts established by the components, which, in turn, alter the ability to interact with the surrounding environment and the resulting functionality ^5, 9, 10, 18, 19^. The disturbance of the contact network can impede the correct function of a ribonucleoprotein complex and trigger disease ^18, 20, 21^. While in physiological conditions interactions between proteins are tightly regulated ^22^, aberrant a occurs in pathologies such as Amyotrophic Lateral Sclerosis (ALS) that is characterised by aggregation Fused in Sarcoma Fus and its partners ^23^.

Multivalency ^11, 12, 13^ is required to keep the dynamicity of the macromolecular complex and is controlled through multiple binding sites ^12^ that are either in structural domains or intrinsically disordered regions (IDRs). The IDRs can contain weak adhesive elements (high stickiness / reactivity to bind) that allow the interaction with multiple partners (promiscuity), including the molecule itself ^24, 25^. This property is the hallmark of Prion Like Domains (PrLD) that are able to favour nucleation of large macromolecular assemblies ^26, 27, 28, 29, 30^. Also RNA molecules have been also reported to modulate the dynamicity ^31, 32^ and the phase separation process of biomolecular condensates ^33, 34, 35, 36^. Indeed, RNAs are flexible and low-complexity polymers with intrinsic high multivalent capacity to interact with several RNA binding proteins (RBPs) and domains (RBDs). Specific RNA interactions result in abrogation of aggregation ^37^ and solubilisation of proteins ^31^ or, by contrast, increased condensation ^20^.

The mechanisms through which proteins phase separate are at present unclear ^8^. With the aim of shedding light on the formation of biomolecular condensates, we computationally analysed the phase separation propensity of *S. Cerevisiae* and *H. sapiens* proteomes. We found that both RBDs and PrLDs co-occur in proteins that are highly prone to phase separate into condensates. Using the SG protein Pub1, we experimentally validated our computational analyses and investigated the effects of RBDs and PrLDs on the formation of condensates. By monitoring Pub1 assembly and its capacity to disturb cell homeostasis, we found that the prion domain leads the assembly process whereas the RNA regulates the dynamicity of the condensate.

## Results

### Phase separation propensity is encoded in both RBDs and PrLDs

To investigate the biophysical proprieties that determine the ability of proteins to phase separate, we first analysed the composition of the best known membrane-less organelle, the SG ^8, 16^ (**Figure 1A**). With respect to the *S. cerevisiae* proteome, we observed a significant enrichment of proteins classified as RBPs ^16^ and prions ^29^ in SGs (p-value <0.001; Fisher’s exact test; **Figure 1A**). This finding suggests that prion and RNA-binding domains are characteristic features of phase-separating proteins, and agrees with previous studies in which lack of structure and nucleic-acid binding abilities were reported to be enriched in SGs ^38, 39, 40^.

**Figure 1.**
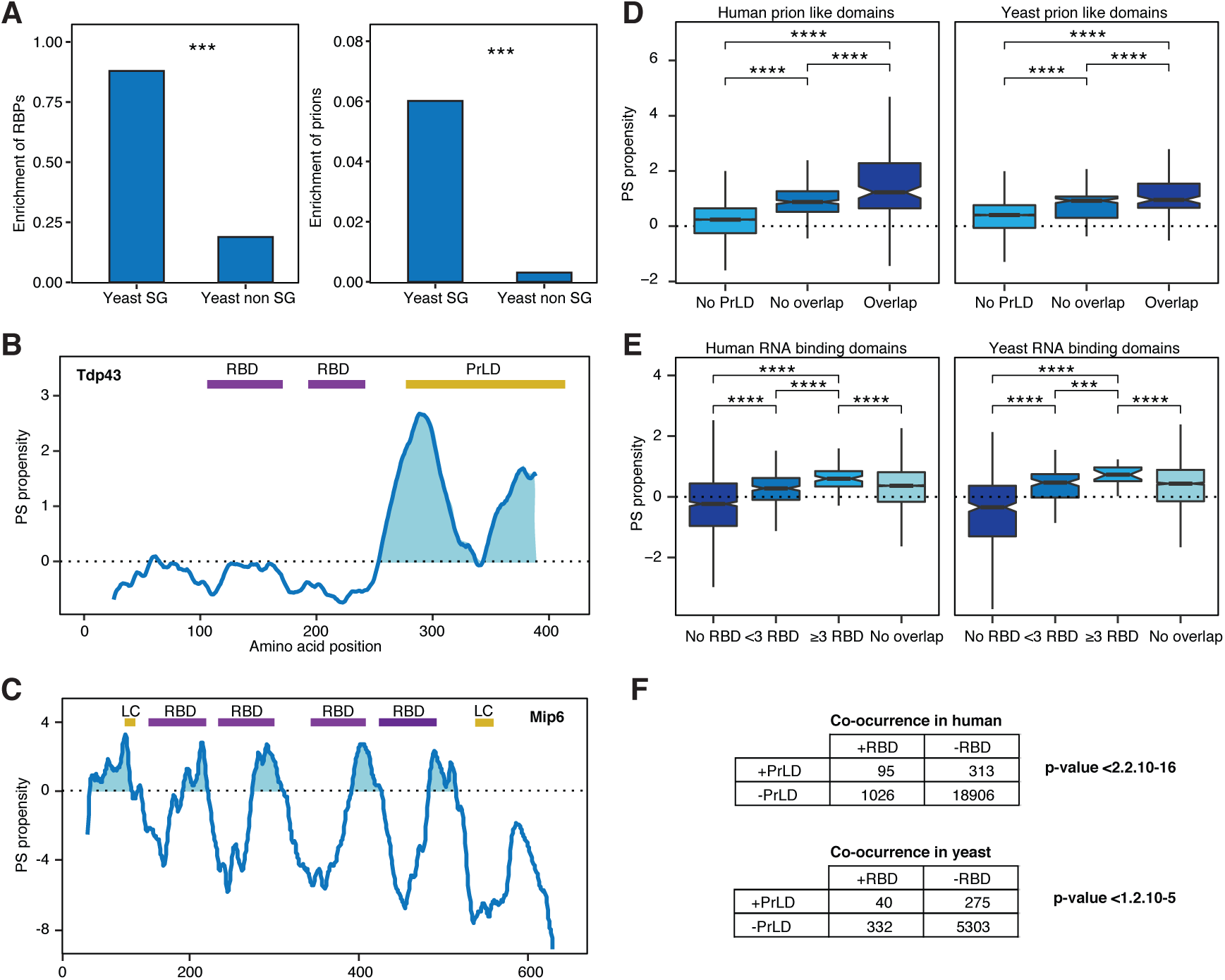
RBDs and prions are associated with the proteins phase separation ability. **A**) Stress granules are enriched in RNA-binding proteins (RBPs) and prions. Left, percentage of RBPs^49^ found in the yeast SGs ^16^ and in the rest of the proteome ^50^. Right, percentage of prion proteins^29^ in same datasets. ***: p-value < 0.001; Fisher’s exact test. **B**) catGRANULE phase separation (PS) propensity profile predicted for Tdp43. The figure also shows two RBDs (Pfam annotated) in purple and a PrLD (PLAAC algorithm prediction) in yellow ^46^. **C**) catGRANULE phase separation (PS) propensity profile predicted for protein Mip6 from S. cerevisiae. The figure shows 4 RBDs in purple and two low-complexity (LC) regions in yellow, as annotated at the ELM database ^51^. The two RBDs close to the N-terminus are essential for the phase separation ^40^. **D**) Box plots showing the percentage of overlapping between PrLDs and the highest peak of phase separation predicted by catGRANULE. Overlap: ≥20% PrLD propensity predicted using PLAAC (PLAAC peak ^46^); No overlap: <20%; No PrLD: PLAAC does not predict PrLD. **E**) Box plots showing the percentage of overlapping between RBDs (Pfam annotated 30357350, scanned with HMMER 29905871) and the highest peak of phase separation predicted by catGRANULE. The proteins have been classified accordingly to the number of RBDs contained in their sequence: < 3 RBD, fewer than 3; ≥ 3 RBD, 3 or more; 0 RBDS, no Pfam annotation. No overlap indicates <20% of sequence coincidence between an RBD and the and the highest peak of phase separation. For all box plots: box represents interquartile range (IQR); central line, median; notches, 95% confidence interval; whiskers, 1.5 times the IQR. ****: p-value <0.0001; t-test. **F**) Fisher test of the association between PrLDs and RBDs. Proteome datasets interrogated for presence (+) or absence (-) of PrLDs (predicted with PLAAC) and RBDs (scanned with HMMER, based on Pfam).

Our observations are in accordance with the principles used to build the *cat*GRANULE method that estimates the phase separation (PS) propensity of proteins ^20, 40^ using structural disorder and RNA-binding propensities (**Figure 1B** and **1C**). Two examples of *cat*GRANULE predictions for proteins containing PrLDs and RBDs are (i) TAR DNA binding protein 43 (TDP-43), associated to ALS disease ^41^, and (ii) Mip6, a yeast protein involved in nuclear mRNA export^40^. *cat*GRANULE profile of Tdp43 shows that the PrLD in the C-terminal region overlaps with the main peak of PS propensity (**Figure 1B**). In agreement with this observation, several peptides of the C-terminal fragment of Tdp43 were found to be the principal components of aggregates in brains of ALS patients ^42, 43, 44^ and we validated that the role of the PrLD in promoting aggregation ^45^. Similarly, *cat*GRANULE predicts that the PS propensity of MIP6 is encoded in a low-complexity domain and 4 RBDs (**Figure 1C**). We proved that MIP6 requires the low-complexity domain and two RBDs close to the N-terminus to undergo phase separation ^40^, thus supporting the correctness of our predictions and the importance of multivalence in particular ^11, 12, 13^.

With the aim of assessing the contributions of PrLDs and RBDs to phase separation, we interrogated both the *H. sapiens* and *S. cerevisiae* proteomes and investigated regions that exhibit strong PS propensity (*cat*GRANULE peaks). We found that if the *cat*GRANULE peak overlaps with at least one PrLD ^46^ there is higher PS propensity than in absence of PrLD or overlap (**Figure 1D**). The same analysis between the *cat*GRANULE peak and one RBD ^47, 48^ showed a stronger PS propensity when there is not overlap. However, based on our previous work with Mip6 ^40^ and the RBPs multi-domain character, we repeated the analysis by considering different numbers of RBDs occurring. We observed that the PS propensity increases with the number of RBDs, and the highest *cat*GRANULE scores are with 3 or more domains (**Figure 1E**). This result suggests a threshold of RBDs required to achieve enough PS potential.

Since both RBDs and PrLDs have been found significantly enriched in phase-separating proteins ^19, 20, 40^, we hypothesized that the two domains might be present and perhaps cooperate to induce condensation. For that reason, we tested whether their co-occurrence is significant. To do so, we quantified the proteins that contain both RBDs and PrLDs and compared them to proteins that contain just one of the two domains or that do not have any. We found a significant co-occurrence of prion and RNA-binding domains in both human and yeast proteomes (Fisher’s test; **Figure 1F**).

### A model to monitor changes in phase separation

We proceeded to validate *in vivo* the link between the ability to phase separate and the occurrence of PrLDs and RBDs. We specifically aimed to clarify the role and implications of each domain type in the phase separation process.

Following the results of our computational analysis, we searched for a phase-separating protein candidate containing RBDs and 1 PrLD (**Figure 2A** and **2B**). To this aim, we focused on *S. cerevisiae* that (i) has been widely validated as a model to study phase separation ^12, 19, 52^, (ii) produces abundant prion-like proteins ^29, 46^ and (iii) has membrane-less organelles and protein quality control system conserved with high eukaryotes ^53, 54, 55^.

**Figure 2.**
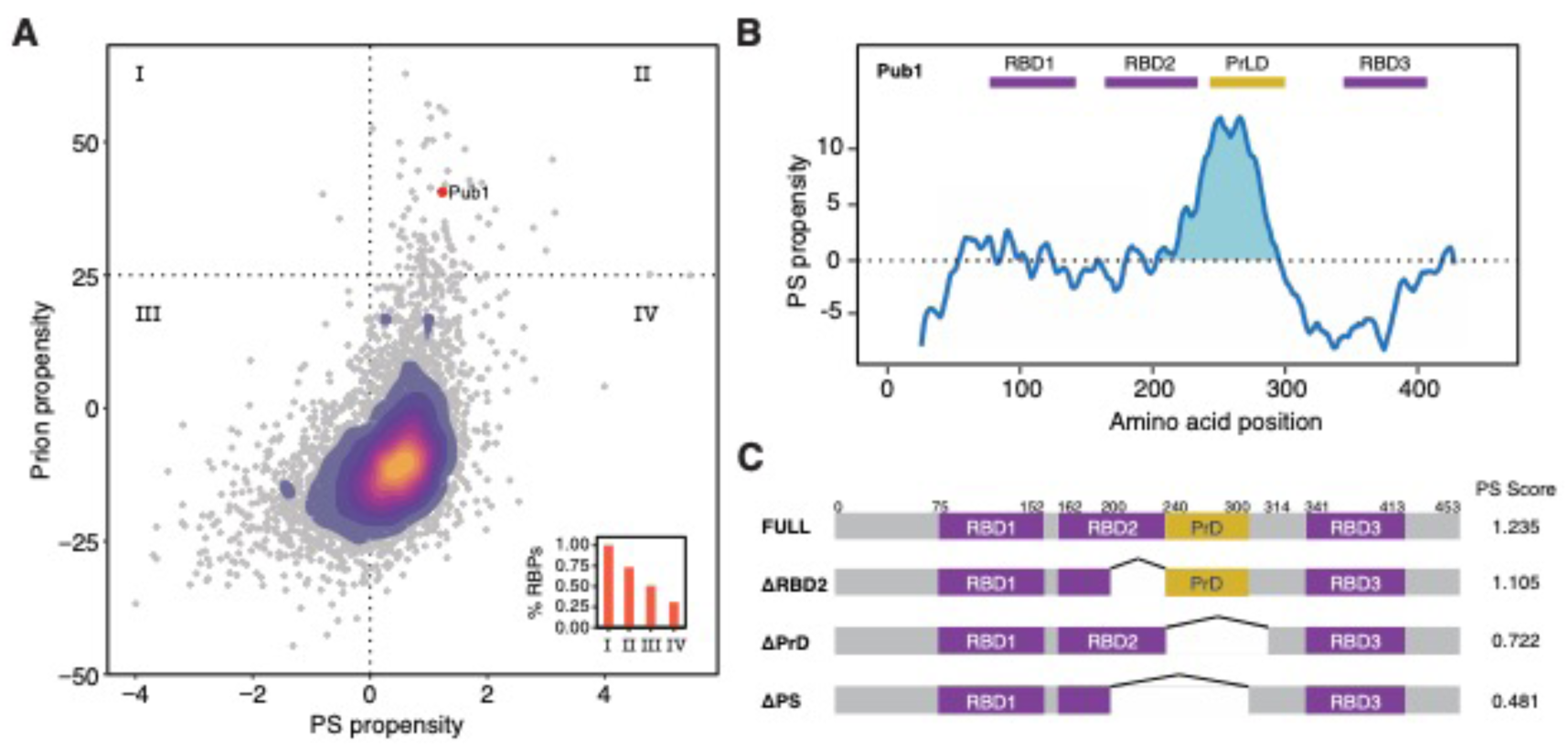
The RBP prion Pub1 model to study phase separation. **A**) Scatter plot and over-imposed 2D-density plot indicating the phase separation (PS) propensity (catGRANULE, X-axis) ^20, 40^ and prion propensity (PLAAC, Y-axis) ^46^ for the yeast proteome. The confidence thresholds of the predictors are highlighted with dotted lines. Inside, bar plot indicating the percentage of RBPs in each of the quadrants. Pub1 protein (with 3 RBDs) is indicated with a red dot. **B**) catGRANULE prediction of Pub1 PS propensity (Y-axis) along sequence (X-axis). The figure also shows 3 RRMs (Pfam annotated, RRM, PF00076 ^47^) in purple and a PrLD (PLAAC prediction) in yellow. **C**) Diagram of the different Pub1 variants studied in this thesis. Targeted deletions of Pub1 full: ΔPrLD (Δ240-314), ΔRRM2 (Δ200-240), ΔPS (Δ200-300).

In order to select a suitable protein candidate, we ranked the whole yeast proteome according to the phase separation (predicted by *cat*GRANULE) ^20, 40^ and the prion propensities (predicted by PLAAC; **Figure 2A**) ^46^. We found 94 proteins containing PrLD (PLAAC score > 25) and prone to phase separate (*cat*GRANULE score > 0): the potential to form prions has been previously investigated experimentally for 55 of them ^29^, 44 are nucleic-acid binding or have annotated RBDs ^56^ and 37 phase separate in fluorescent *foci* ^29^. Only 7 belong to the ≥3 RBDs group, and Pub1 is the only prion with 3 RBDs and a phase separation propensity at the top decile in comparison with the whole proteome (**Table S5 ∼** and **Figure S1**).

The Pub1 protein binds polyuridilated RNA and is present in SGs under different stresses ^19, 57^. The Ps peak ^20, 40^ of Pub1 overlaps with the second RBD (RBD2, a structured RNA Recognition Motif, RRM) and the PrLD ^58^ (**Figure 2B, Figure S2** and **Figure S3**). Thus, in the PS region we have the two domains. To investigate the contributions of both the PrLD and the RBD, we performed deletion of these regions resulting in three constructs (**Figure 2C**): ΔRBD2, lacking the part of the second RRM region that is under the phase separation peak; ΔPrLD, lacking the PrLD; and ΔPS, lacking the overall region under the phase separation peak.

We monitored Pub1 expression and phase separation by fusing the gene to GFP and employing a strong inducible promoter (GAL1). In this way, we ensured enough Pub1 concentration to trigger its accumulation into condensates independently from environmental stresses ^19, 57^. Based on the multivalence of RBDs ^11, 12, 13^ and *cat*GRANULE predictions (**Figure 2C** and **Figure S4**) ^20, 40^, we expected that deletion of specific domains decrease the PS potential.

### PUB1 PrLD guides phase separation

To test the effect of PrLDs and RBDs on Pub1 protein phase separation, we analysed the expression levels of our four Pub1 variants (**Figure S5**) and their localization at different expression times (**Figure 3A**). Induction resulted in similar expression levels (**Figure S5**) and comparable soluble/insoluble fractions (**Figure S6**), but the assemblies formed b Pub1, ΔPrLD ΔRRM2 and, ΔPS and variants looked dramatically different. We analysed and classified cells in 3 groups according to the presence and size of condensates: diffuse (no condensates), small condensates (<1µm) and big condensates (≥1µm) (100 cells were counted for each strain; **Figure 3B**).

**Figure 3.**
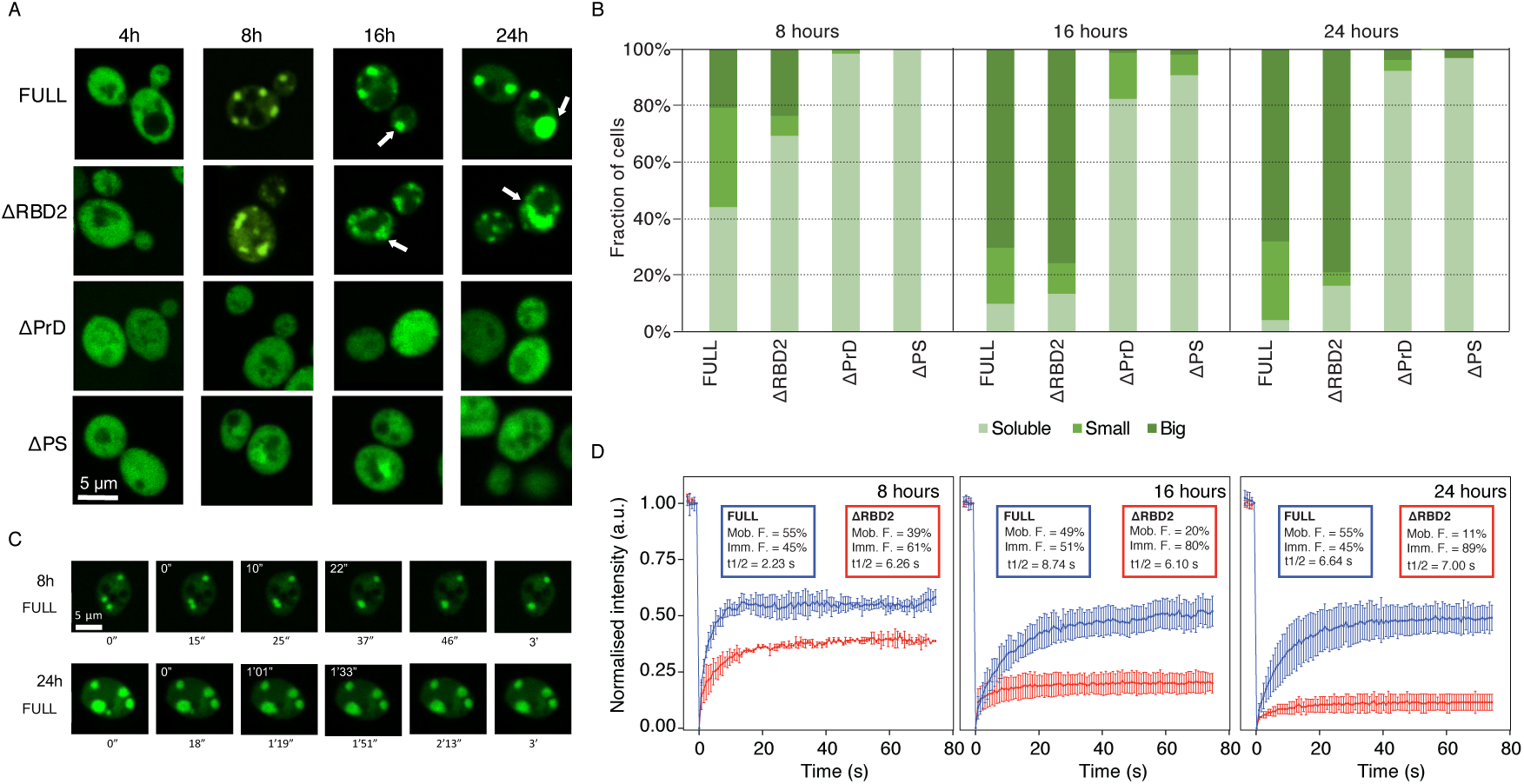
Pub1 expression and localization. **A**) Representative pictures of Pub1 variants provides evidence that the PrLD has a critical role for phase separation. From **top** to **bottom**, Pub1 variants are ranked by decrease in multivalency. From **left** to **right**, time progresses in hours. Arrows indicate round condensates in the FULL variant and less round condensates in the ΔRBD2. **B**) Cells are classified in three phenotypes: Big, cells that present ≥1 µm condensates; Small, cells that present <1 µm condensates; Diffuse: cells that present fluorescent homogenously distributed. Percentage of cells (Y-axis) are normalized to the total number of fluorescent cells. A minimum of 100 cells were counted for each strain and condition. **C**) Fusion events of the condensates formed at different expressions times of Pub1. White numbers indicate the timing of fusion events. **D**) ΔRBD2 and Pub1 Fluoresce Recovery After Photobleaching (FRAP) measurements of the condensates at different expression times. Plots indicate mean and standard deviations of normalized fluorescence over time, background subtracted. Experiments correspond to three biological replicates. Percentages of mobile (Mob. F.) and immobile (Imm. F.) protein fractions. Protein half-life (t1/2) in seconds.

After 4 hours under GAL1 overexpression all strains showed diffuse protein distribution through the cytosol (**Figure 3A**). However, after 8 hours we observed a different condensation: 56% Pub1 and 31% of ΔRBD2 populations presented small or big condensates, whereas ΔPrLD and ΔPS proteins remained mainly diffuse across the populations (**Figure 3B**). The condensates became larger with prolonged induction times (**Figure 3A** and **3B**). After 16 hours of overexpression nearly all Pub1 and ΔRBD2 cells showed condensates and most of them were classified as big (70% and 76%, respectively). In the case of ΔPrLD and ΔPS populations, the number of cells with condensates remained below 18% and 9%, respectively. Importantly, the number of condensates is increased in Pub1 cells and their size is larger in ΔRBD2 cells (**Figure 3A** and **3B**).

### Truncation of RBD2 results in less dynamic condensates

Through microscopy cell screening we found that the number and size of the condensates increase over time (**Figure 3A** and **Figure 3B**). In agreement with these observations, larger and irregular condensates were previously associated with lower dynamicity and aging (e.g. formation of aggregated forms) ^18, 21^.

To test if the increase in size is associated with a change in the dynamicity of the assemblies, we recorded fusion events of Pub1 condensates at different expression times. While at 8 hours we observed fast fusion events (22 seconds) that result in round condensates, after 24 hours the fusion events are 4 times longer (1.33 seconds) and the final condensates do not have a round shape (**Figure 3C**).

Thus, in addition to a size increase, there was a drastic change in shape (**Figure 3C**). The images indicated that ΔRBD2 condensates look more irregular than those with Pub1 (**Figure 3A** and **3B**), suggesting condensates with different material states. To confirm if the change is associated to a decrease in dynamicity, we used Fluoresce Recovery After Photobleaching (FRAP) measurements, at different expression times, on both Pub1 and ΔRBD2 condensates and measured their diffusion properties (**Figure 3D** and **Figure S7**). We observed that Pub1 condensates exhibit similar recovery at different induction times (50-60%). By contrast, ΔRBD2 presents slower recovery that further decreases after long induction times (reaching less than 10% after 24h of expression). This finding reveals that condensates formed with ΔRBD2 have more solid-like character than those formed by Pub1.

### Phase separation propensity correlates with cell growth impairment

We found that Pub1 variants differ in PS propensity and dynamicity (**Figure 3C** and **3D**), which indicates a change in their capacity to interact with the surrounding environment. Indeed, the ability of molecules to interact with each other has a strong effect cellular homeostasis ^59^ and tight regulation is required to avoid damage ^60^.

We employed a growth assay to measure how Pub1 variants disturb cell homeostasis. Under GAL1 promoter, PUB1 expression is controlled by the carbon source present in the media: i) glucose acts as an inhibitor allowing growth without Pub1 expression or (after a media change) stopping its expression; ii) galactose acts as an inductor allowing growth when Pub1 is expressed (**Figure 4A**). At no-inducing conditions all strains grow similarly, both in liquid (**Figure 4A**) and solid media (**Figure S8**), as expected given their isogenic origin and the absence of metabolic differences before induction. Overexpression of eGFP alone results in no significant decrease in growth rate, supporting yeast robustness against proteotoxicity (**Figure S8**). However, the overexpression of Pub1 variants results in a dual phenotype:ΔPS and ΔPrLD principally diffuse while ΔRBD2 and PUB1 condense (**Figure 3**) and their growth curves show gradual range of doubling times (**Figure 4B-C, Figure S8, Table S1**). Importantly, we observed a decrease in doubling time associated with Pub1 expression that correlates with the phase separation potential of each variant (**Figure 4D**).

**Figure 4.**
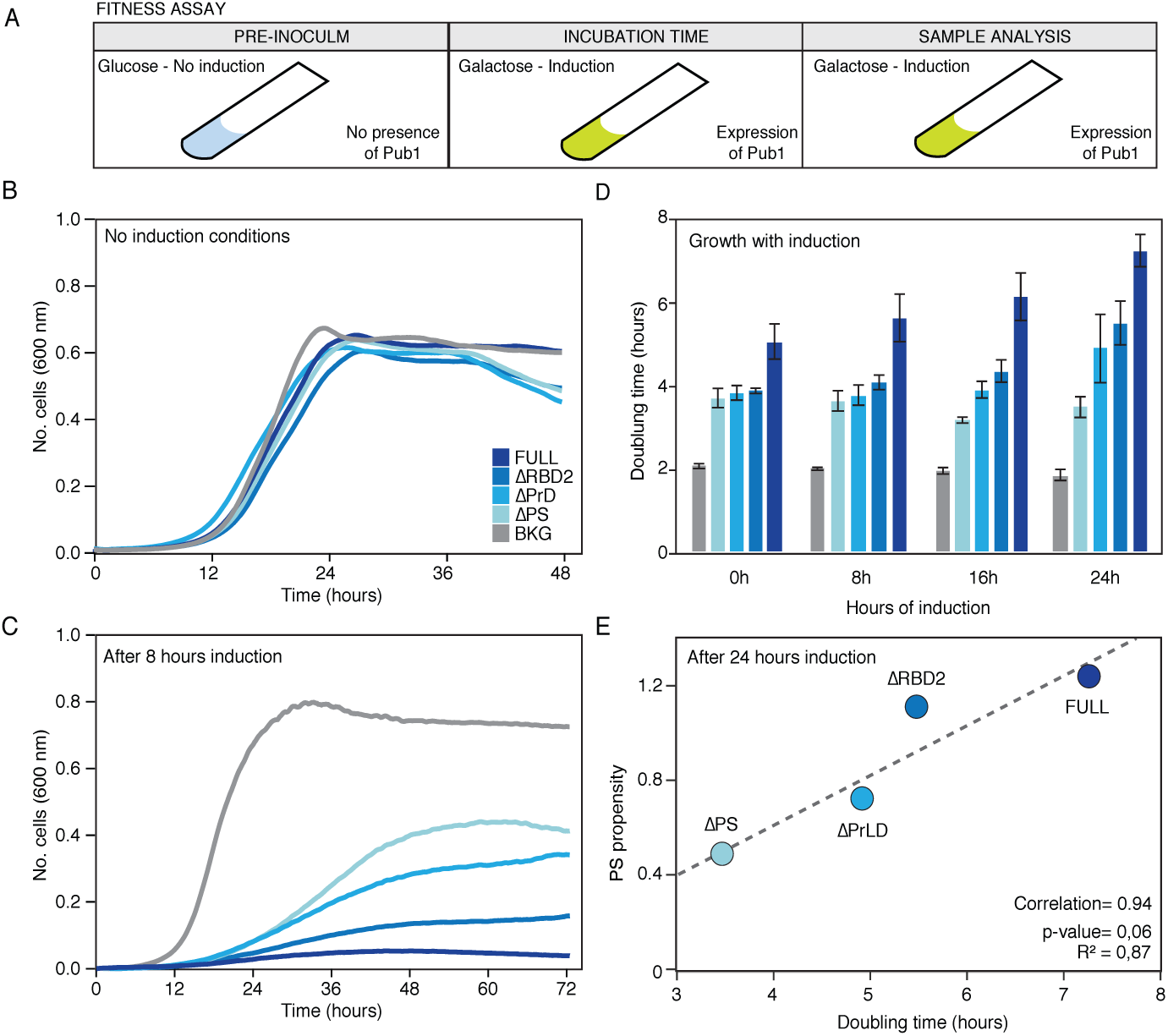
Changes in the growth curve after the expression of the different Pub1 variants. **A**) Scheme of the different steps followed to prepare the cell samples before the growth curve analysis. The pre-inoculum was in glucose media, then the cells were incubated in inducing conditions during different times (0, 8, 16 and 24 hours; **Supplementary Table S1** and **S2**) before growth characterization in presence of galactose (the inductor). **B)** Growth curves in no-inducing conditions (glucose) to control background fitness between strains. **C**) Growth curves where monitored for three days in inducing conditions (see **Supplementary Material Tables S1**). The graph exhibits growth curves in presence of inductor and after 8 hours of pre-induction. **D**) Bar plots indicating the doubling time of different variants measured after pre-inducing times. For lag time and saturation limit see **Supplementary Material Table S1. E**) Linear correlation between the propensity to phase separation calculated with catGRANULE and the doubling time measured after 24 hours of induction.

Flow cytometry experiments indicate that the growth differences associated with our variants are not caused by plasmid loss or cell death (**Figure S9, S10 and S11** and **Table S2**) but a perturbation in the cell division process. The expression of Pub1 always exhibits a larger growth impairment, and after 24 hours its doubling time is 1.44 times slower than when the induction was started (**Figure 4C** and **Table S1**). ΔPrLD and ΔRBD2 doubling times also gradually decreased with the induction time, whereas ΔPS growth speed remains constant. We found that the *cat*GRANULE score correlates with the doubling time (**Figure 4E**), which suggests an connection between impairment of cell division and aberrant formation of phase-separated assemblies. In summary, our results indicate that the propensity of Pub1 to phase separate into specific species, and not just condensation *per se* ^59^, is associated with the ability to disturb cell growth.

### Phase separation propensity correlates with fitness recovery time

To further investigate how condensates with different dynamicity affect cellular functioning, we measured the cell capacity to reacquire physiological conditions after Pub1 variants overexpression. This assay is intrinsically linked to the diffusion capacities of Pub1 and its variants and thus the reversibility of the condensate state ^19, 61^. In these experiments, after different induction times, we moved the strains to glucose that inhibits the galactose pathway and thus expression of PUB1.

After the expression of PUB1 variants is blocked (**Figure 5A**), all the strains showed growth recovery (**Figure 5B-C, Figure S12** and **Table S3**), however the effect decreases with induction time (**Figure 5D**). This agrees with the growth impairment effect that we previously found associated with the induction time (**Figure 4A** and **Figure S13**). For the different variants, we observed a consistent change in number, size and dynamicity (FRAP) of condensates linked to the induction times. Interestingly, a similar decrease in recovery capacity has been associated with “aging” in the case of FUS condensates, whose loss of dynamicity and progressing aggregation was recently investigated *in vitro* ^18, 20, 21^.

**Figure 5.**
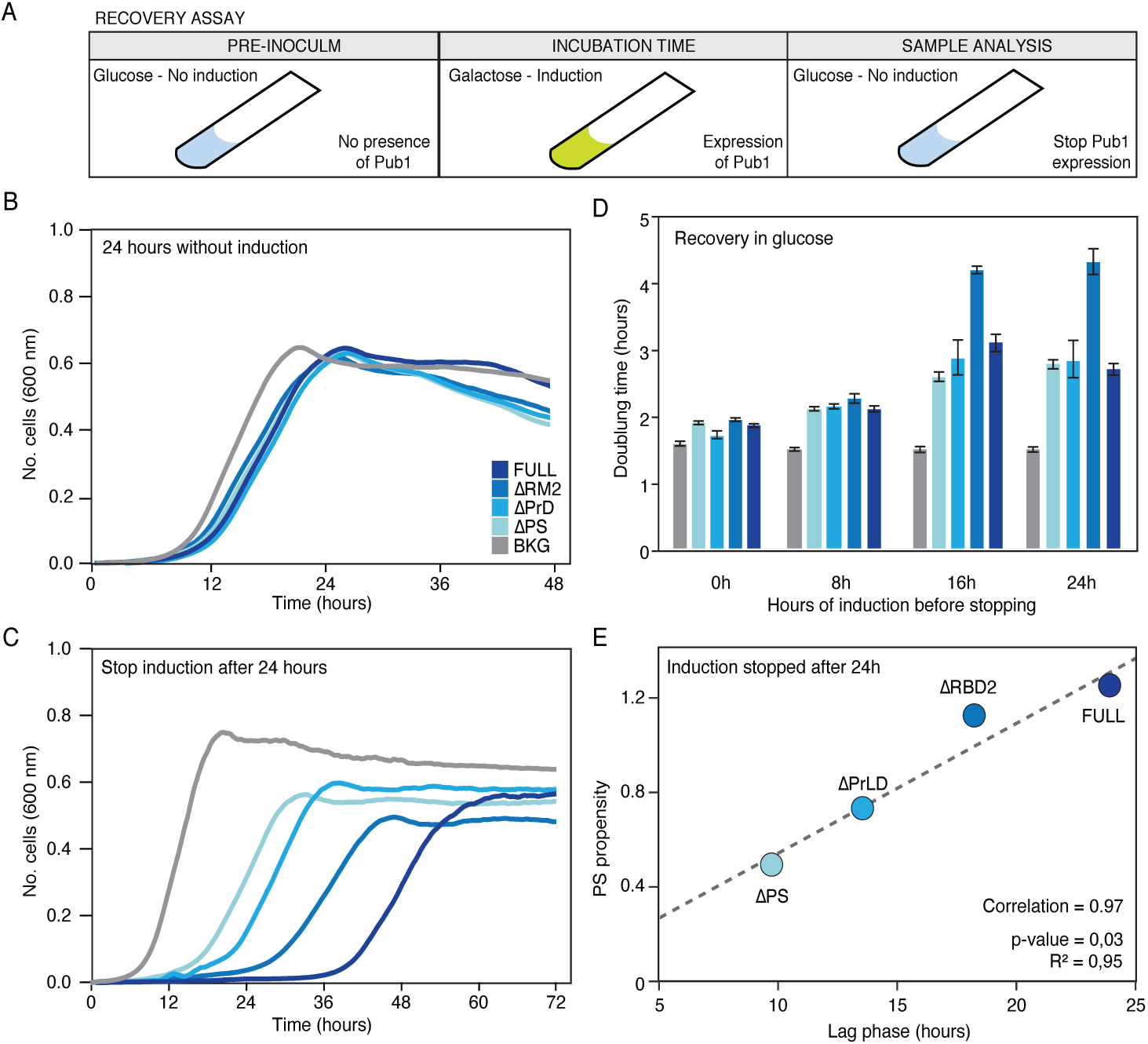
Changes in the growth recovery after the expression of the different Pub1 variants. **A**) Scheme of the different steps followed to prepare the cell samples before the growth curve analysis. The pre-inoculum was in glucose media. Then the cells were incubated in inducing conditions during different times (0, 8, 16 and 24 hours) before changing again to the glucose media (inhibition conditions) to stop Pub1 expression and perform the growth characterization. **B**) Growth curves after 24 hours growing in no-inducing conditions as a fitness control. **C**). Growth curves where monitored for three days after stopping PUB1 expression (see **Supplementary Material Table S3**). The graph exhibits growth curves after 24 hours of pre-induction. For other pre-induction times see **Supplementary Material Table S3. D**) Bar plots indicating the doubling time once Pub1 expression is stopped and after different pre-inducing times. For lag time and saturation limit see **Supplementary Material Table S3**. Bar high corresponds to arithmetic mean of three independent biological replicates and error bars indicate standard deviations. **E**) Linear correlation between the propensity to phase separate calculated by catGRANULE and the lag time measured after 24 hours of induction

Focusing on the growth curve parameters, we observed that, for a specific induction time, the doubling time and saturation level remain quite similar between strains, whereas the lag time is different and fits, again, with the growth impairment previously measured (**Figure S13**). It should be mentioned that in a growth curve, lag time informs about the time that a population of cells requires to achieve the top division speed. Thus in our case, after stopping induction, the lag time informs about the time required by the cell to overcome the disturbance caused by the overexpressed Pub1. Importantly, the lag time is the parameter that best correlates with the phase separation propensity of Pub1 variants (**Figure 5E**).

At long induction times, we observed a strong slow-down of ΔRBD2 doubling time. Whereas the growth rate recovery obtained after 24 hours of induction is around 3 hours for Pub1, ΔPS and ΔPrLD, division of ΔRBD2 cells takes more than 4 hours. For nearly all the fitness analyses ΔRBD2 and Pub1 were close related, with ΔRBD2 performances slightly better than Pub1 (**Figure 3A** and **3C** and **Figure 4C** and **4D**). Yet, ΔRBD2 condensates are less dynamic (**Figure 3D**), a characteristic that that is associated with lower reversibility and more difficult disassembly ^18, 20, 21^. In the future, we plan to investigate more these aspects, although we can already hypothesize that the lower dynamicity of ΔRBD2 condensates has a deep effect on cell fitness.

### PUB1 PrLD interacts with numerous proteins with essential cellular functions

PrLD domain is crucial to achieve the formation of condensates (**Figure 2**) and PS propensity is intimately associated with the capacity to disrupt cell function (**Figure 4**). Upon PrLD depletion, we found a fast recovery of cell fitness. Thus, PrLD cause cell toxicity by inducing condensation and interacting with the surrounding environment through. To shed light on this, we proceeded to measure changes in Pub1 protein network upon PrLD (ΔPrLD) and phase separation (ΔPS) peak depletions.

Protein networks were studied using immuno-precipitation (**Figure S14** and **S15**) and mass-spectrometry (**Figures S16-S19** and **Table S4**). Specifically, we conducted our experiments after 8 hours of induction, when the percentage of fluorescent cells is close to 50% (**Figure S10** and **Table S2**) and the differences between strains are measurable (**Figure 1** and **Figure 4**).

We found a dramatically different number of Pub1, ΔPrLD and ΔPS interactors: 436, 49 and 23, respectively (false discovery, FDR, of 0.001; **Figure 6A-B** and **Table S4**). When compared to the rest of the proteome (around 4000 proteins ^50^), Pub1 and ΔPrLD interactors showed more structurally disordered and nucleic acid binding proteins (as predicted by the *multi*CM algorithm that calculates enrichments in physico-chemical properties; see **Supplementary Material; Figure 6C**) ^62^. By contrast, ΔPS interactors are less prone to bind to nucleic acids and have a propensity to undergo solid aggregation similar to the rest of the proteome (**Figure 6C**) ^63^. Proteins bond by ΔPrLD and ΔPS are more hydrophobic and less nucleic acid binding than Pub1(**Figure 6D**).

**Figure 6.**
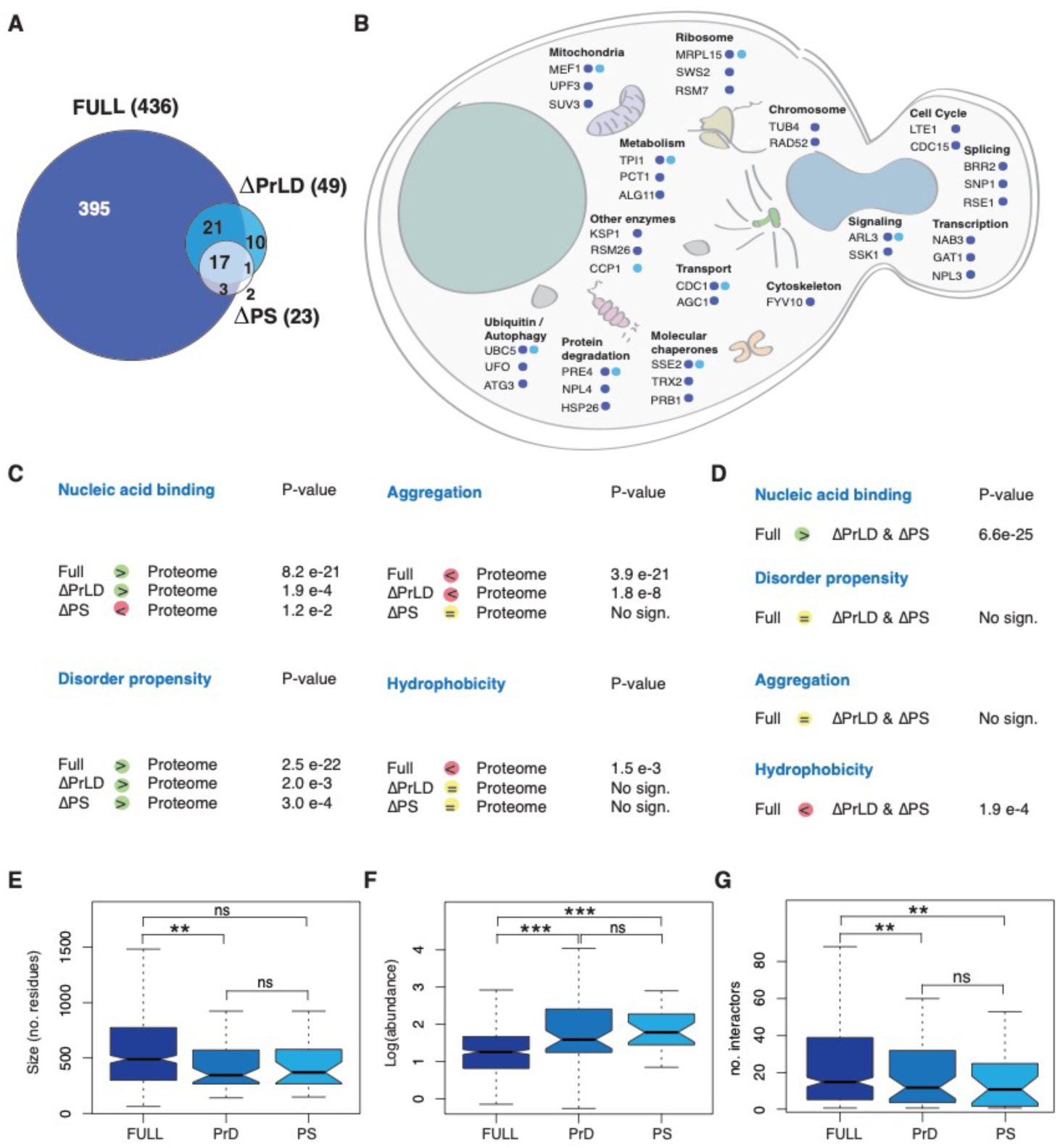
Differences between the interactors of the Pub1 variants. A) Venn diagram represents the number of interactors of each protein variant in the conditions of the experiment (after 8 hours of overexpression). FDR = 0.001. B) Representation of Pub1 variants interactors in the context of a yeast cell. Representation of the different sequestered proteins accordingly to the cellular process and location. C) Comparison of properties between Pub1 variants interactors and SG proteins or proteome accordingly to multiCM. D) Comparison of properties among Pub1 variants interactors accordingly to multiCM. E) Boxplots showing the size distribution of the interactors. F) Boxplots showing the abundance distribution of the interactors (PaxDB: GPM, Aug 2014). G) Boxplots showing the interaction capacity of the interactors (BioGRID). For all box plots: box represents interquartile range (IQR); central line, median; notches, 95% confidence interval; whiskers, 1.5 times the IQR. **:p-value<0.05, ***: p-value <0.005 (Mann Whitney).

Moreover, Pub1 interactors are longer (**Figure 6E**), less abundant (**Figure 6F**) and have a larger number of interactors (**Figure 6G**) than proteins contacted by ΔPrLD and ΔPS (PaxDB ^64^ and BioGRID ^65^ were used for this analysis), which is compatible with the observation that expression of genes with a large number of partners is tightly controlled to avoid massive aggregation in the cell ^66, 67^. In the list of Pub1 interactors, we counted 106 (24%) proteins with strong-confidence PS propensity (*cat*GRANULE score > 1), of which 16 are experimentally-validated SG (**Table S7**) proteins ^16^. Interestingly, all the characteristics associated to phase-separation (long, RNA-binding and highly interacting) ^20, 68^ are intimately connected with the presence of PrLD in Pub1.

The PrLD is essential for the recruitment of Pub1 interactors: 387 Pub1 partners (>90%) are not present in ΔPrLD network and 32 of ΔPrLD (>60%) interactors are not present in ΔPS network (**Figure 6A**). Additionally, presence of PrLD in Pub1 is associated with recruitment of 22 PrLD-containing proteins (**Table S7**), of which 15 have been previously reported to bind Pub1 (e.g. polyadenylate-binding protein, Pab1; nuclear polyadenylated RNA-binding proteins Nab2, Nab3, Nab6; and mRNA-binding proteins Puf2, Puf3, Puf4; BioGRID ^65^; **Table S7**).

Many different cellular circuits are affected by Pub1 condensation, including transcription (e.g., Nuclear polyadenylated RNA-binding protein 3 Nab3 and Transcriptional regulatory protein Gat1), chromosome organisation (Tubulin gamma chain Tub4 and DNA repair and recombination protein rad52), ribosome and mitochondrion (37S ribosomal protein SWS2 and 37S ribosomal protein S7 Rsm7), degradation and autophagy (Nuclear protein localization protein 4 Npl4, Autophagy-related protein 3 Atg3 and Ubiquitin ligase complex F-box protein Ufo1) and molecular chaperones (Heat shock protein homolog SSE2 and prion Cerevisin Prb1). The number of proteins that we detected is proportional to the PS propensities of Pub1 variants (**Figure 6B** and **Figure S19**). Removal of PrLD results in depletion of proteins associated with cell cycle, transcription, translation and lipid metabolism. Similarly, the PS peak is associated with ribosome, spliceosome, transport and mitochondrial biogenesis.

To understand to what extent alteration of protein abundance affects cell fitness, we analysed the amount of essential and dosage sensitive proteins (**Table S7**) sequestered by Pub1. We obtained that Pub1 interacts with 65 essential (14% of the whole interactome, including Pav1, Cdc15, Tub4 and Taf11) and 92 dosage sensitive proteins (20%, including Hsp82, Gbp2, Pub1 and If4f1).

In summary, our analysis indicates that Pub1 interacts with proteins involved in many functional pathways. Loss of these interactions result in a disruption of cell homeostasis, which causes impediments in cell growth. Toxicity is intimately linked to the amount of proteins sequestered by Pub1 condensates and by their ability to participate in large interaction networks.

## Discussion

There is an intense debate on which protein regions contribute to the assembly and dynamics of biomolecular condensates ^30, 31, 37, 69, 70, 71, 72^. Our computational analysis of *H. sapiens* and *S. cerevisiae* proteomes reveals that the PS propensity is intrinsically linked to occurrence of PrLDs and RBDs. We used the SG protein Pub1 to investigate the specific contributions of these domains designing mutations with the *cat*GRANULE approach ^40^.

Our results clearly show that PrLD is required for condensation and RBD modulates its dynamics. Removal of the RBD reduces the liquid-like behaviour of the condensate ^22^ in favour of interactions that lead to more solid-like aggregation ^29, 73^. In agreement with this finding, previous studies indicated the importance of RNA in SGs network specificity and organization ^74^: RNA-binding proteins can be dragged in the condensates by RNA networks ^31, 37, 75^. Indeed, RNA is a key modulator of the dynamics and material state of ribonucleoprotein complexes ^70, 76, 77, 78^ and a highly structured RNA can provide the scaffold to allow formation of large assemblies ^20, 31^. For instance, the long non coding RNA NEAT1 promotes condensation of nuclear paraspeckle components ^71, 79, 80^. In many other cases long RNAs could be able to recruit RBPs, easily achieving the critical concentration for phase separation ^20, 75, 81^.

In line with previous reports ^9, 16^, our analysis of protein interactions indicates that other PrLDs are recruited into condensates. Thus, PrLD interactions could be essential to seed condensates formation ^82^ and once a critical mass of interactions is reached, the condensate would gather additional molecules establishing a large network of protein-protein (PPIs) and protein-RNA (PRIs) interactions ^68^. One could also hypothesize the intriguing possibility that PrLDs might establish weak interactions with nucleic acids to promote condensation. Indeed, disordered regions containing PrLDs have been shown to have some propensity to bind RNA ^49, 83^.

‘Is condensation the cause or rather consequence of cell toxicity? Heterologous over-expression of proteins, such as for instance human Tdp43, causes toxicity in *S. cerevisiae* because, due to the absence of a functional homologue, the protein establishes aberrant interactions with other molecules disturbing cell homeostasis ^45^. Aggregation in solid-like aggregates represents a way to reduce the interaction potential of Tdp43 and its consequent damages to the cell ^84^. By contrast, autologous over-expression of proteins, such as for instance Pub1, is linked to other effects. In this case toxicity arises for i) the imbalanced complex stoichiometry caused by over-expression and ii) the subsequent formation of assemblies ^85^ in which Pub1 and its interactors become unavailable to perform their normal physiological functions. Thus, aberrant condensation creates the conditions for inappropriate molecular sequestration (loss of function) ^86^ and occurrence of undesired reactions (gain of function) ^87^.

Protein interactors of the different Pub1 variants are related to transcription, translation, folding and degradation. The variety of processes affected by condensations points to a general disruption of homeostasis. Indeed, Pub1 interactors are enriched in essential and dosage sensitive proteins. Importantly, the presence of the PrLD not only favours the recruitment of more proteins, but also highly interacting proteins, which may have an amplification effect on the final network size. As suggested by our mass spectroscopy results, the main cause of the fitness decrease could be deficient cell division. Indeed, Pub1 condensates interact with proteins related to cell cycle arrest, whereas the less toxic variants, ΔPrLD and ΔPS, do not. We suspect that toxicity arise when the protein interacts with essential cellular machinery, ultimately trapped in condensates.

In conclusion, our results demonstrate that PrLDs and RBDs play different but not independent roles in phase separation. The two domain types have intimately inter-connected tasks: PrLD has a clear role in creating protein interactions and assembling condensates, whereas RDB influences the final material state. The fine interplay between RBDs and PrLDs regulates the formation of membrane-less organelles, inducing quick formation of ribonucloprotein assemblies and promoting their fast disaggregation.

## Contributions

NLG conducted the experiments supervised by NSDG and GGT. AA did the computational analysis together with NSDG and GGT. GC and RMV carried out the mass-spectrometry experiments and their analyses. NLG, NDSG and GGT wrote the paper.

## Acknowledgements

We thank all members of the Tartaglia’s, Vabulas’ laboratory, Dr. Teresa Botta-Orfila and Dr. Benedetta Bolognesi. The research leading to these results has been supported by European Research Council (RIBOMYLOME_309545 to GGT, ASTRA 855923 to GGT and METAMETA_311522 to RMV), Spanish Ministry of Economy and Competitiveness (BFU2014-55054-P and BFU2017-86970-P), ‘Fundació La Marató de TV3’ (PI043296) and the collaboration with Peter St. George-Hyslop financed by the Wellcome Trust. We acknowledge support of the Spanish Ministry of Economy and Competitiveness, ‘Centro de Excelencia Severo Ochoa 2013-2017’. We also acknowledge the support of the CERCA Programme/Generalitat de Catalunya and of Spanish Ministry for Science and Competitiveness (MINECO) to the EMBL partnership.

## References

1. Cirillo D, Marchese D, Agostini F, Livi CM, Botta-Orfila T, Tartaglia GG. Constitutive patterns of gene expression regulated by RNA-binding proteins. Genome Biol 15, R13 (2014).

2. Stockmayer WH. Molecular distribution in condensation polymers. Journal of Polymer Science 9, 69–71 (1952).

3. Banani SF, Lee HO, Hyman AA, Rosen MK. Biomolecular condensates: organizers of cellular biochemistry. Nat Rev Mol Cell Biol 18, 285–298 (2017).

4. Brangwynne CP, et al. Germline P granules are liquid droplets that localize by controlled dissolution/condensation. Science 324, 1729–1732 (2009).

5. Alberti S. The wisdom of crowds: regulating cell function through condensed states of living matter. J Cell Sci 130, 2789–2796 (2017).

6. Shin Y, Brangwynne CP. Liquid phase condensation in cell physiology and disease. Science 357, (2017).

7. Marchese D, de Groot NS, Lorenzo Gotor N, Livi CM, Tartaglia GG. Advances in the characterization of RNA-binding proteins. Wiley Interdiscip Rev RNA 7, 793–810 (2016).

8. Mitrea DM, Kriwacki RW. Phase separation in biology; functional organization of a higher order. Cell Commun Signal 14, 1 (2016).

9. Markmiller S, et al. Context-Dependent and Disease-Specific Diversity in Protein Interactions within Stress Granules. Cell 172, 590–604 e513 (2018).

10. Alberti S, Gladfelter A, Mittag T. Considerations and Challenges in Studying Liquid-Liquid Phase Separation and Biomolecular Condensates. Cell 176, 419–434 (2019).

11. Boeynaems S, et al. Protein Phase Separation: A New Phase in Cell Biology. Trends Cell Biol 28, 420–435 (2018).

12. Nakamura H, et al. Intracellular production of hydrogels and synthetic RNA granules by multivalent molecular interactions. Nat Mater 17, 79–89 (2018).

13. Li P, et al. Phase transitions in the assembly of multivalent signalling proteins. Nature 483, 336–340 (2012).

14. Brangwynne CP, Mitchison TJ, Hyman AA. Active liquid-like behavior of nucleoli determines their size and shape in Xenopus laevis oocytes. Proc Natl Acad Sci U S A 108, 4334–4339 (2011).

15. Feric M, et al. Coexisting Liquid Phases Underlie Nucleolar Subcompartments. Cell 165, 1686–1697 (2016).

16. Jain S, Wheeler JR, Walters RW, Agrawal A, Barsic A, Parker R. ATPase-Modulated Stress Granules Contain a Diverse Proteome and Substructure. Cell 164, 487–498 (2016).

17. Das R, et al. New roles for the de-ubiquitylating enzyme OTUD4 in an RNA-protein network and RNA granules. J Cell Sci 132, (2019).

18. Alberti S, Hyman AA. Are aberrant phase transitions a driver of cellular aging? Bioessays 38, 959–968 (2016).

19. Kroschwald S, et al. Different Material States of Pub1 Condensates Define Distinct Modes of Stress Adaptation and Recovery. Cell Rep 23, 3327–3339 (2018).

20. Cid-Samper F, et al. An Integrative Study of Protein-RNA Condensates Identifies Scaffolding RNAs and Reveals Players in Fragile X-Associated Tremor/Ataxia Syndrome. Cell reports 25, 3422–3434 e3427 (2018).

21. Patel A, et al. A Liquid-to-Solid Phase Transition of the ALS Protein FUS Accelerated by Disease Mutation. Cell 162, 1066–1077 (2015).

22. Mann JR, et al. RNA Binding Antagonizes Neurotoxic Phase Transitions of TDP-43. Neuron 102, 321–338 e328 (2019).

23. Lenzi J, et al. ALS mutant FUS proteins are recruited into stress granules in induced pluripotent stem cell-derived motoneurons. Dis Model Mech 8, 755–766 (2015).

24. Quiroz FG, Chilkoti A. Sequence heuristics to encode phase behaviour in intrinsically disordered protein polymers. Nat Mater 14, 1164–1171 (2015).

25. Romero P, Obradovic Z, Li X, Garner EC, Brown CJ, Dunker AK. Sequence complexity of disordered protein. Proteins 42, 38–48 (2001).

26. Riemschoss K, et al. Fibril-induced glutamine-/asparagine-rich prions recruit stress granule proteins in mammalian cells. Life Sci Alliance 2, (2019).

27. Sabate R, Rousseau F, Schymkowitz J, Ventura S. What makes a protein sequence a prion? PLoS Comput Biol 11, e1004013 (2015).

28. Cushman M, Johnson BS, King OD, Gitler AD, Shorter J. Prion-like disorders: blurring the divide between transmissibility and infectivity. J Cell Sci 123, 1191–1201 (2010).

29. Alberti S, Halfmann R, King O, Kapila A, Lindquist S. A systematic survey identifies prions and illuminates sequence features of prionogenic proteins. Cell 137, 146–158 (2009).

30. Protter DSW, et al. Intrinsically Disordered Regions Can Contribute Promiscuous Interactions to RNP Granule Assembly. Cell Rep 22, 1401–1412 (2018).

31. Sanchez de Groot N, et al. RNA structure drives interaction with proteins. Nat Commun 10, 3246 (2019).

32. Schuster BS, et al. Controllable protein phase separation and modular recruitment to form responsive membraneless organelles. Nat Commun 9, 2985 (2018).

33. Zhang H, et al. RNA Controls PolyQ Protein Phase Transitions. Mol Cell 60, 220–230 (2015).

34. Berry J, Weber SC, Vaidya N, Haataja M, Brangwynne CP. RNA transcription modulates phase transition-driven nuclear body assembly. Proc Natl Acad Sci U S A 112, E5237–5245 (2015).

35. Shevtsov SP, Dundr M. Nucleation of nuclear bodies by RNA. Nat Cell Biol 13, 167–173 (2011).

36. Mao YS, Sunwoo H, Zhang B, Spector DL. Direct visualization of the co-transcriptional assembly of a nuclear body by noncoding RNAs. Nat Cell Biol 13, 95–101 (2011).

37. Zacco E, et al. RNA as a key factor in driving or preventing self-assembly of the TAR DNA-binding protein 43. J Mol Biol 431, 1671–1688 (2019).

38. Kaganovich D, Kopito R, Frydman J. Misfolded proteins partition between two distinct quality control compartments. Nature 454, 1088–1095 (2008).

39. Kato M, et al. Cell-free formation of RNA granules: low complexity sequence domains form dynamic fibers within hydrogels. Cell 149, 753–767 (2012).

40. Bolognesi B, et al. A Concentration-Dependent Liquid Phase Separation Can Cause Toxicity upon Increased Protein Expression. Cell reports 16, 222–231 (2016).

41. Prasad A, Bharathi V, Sivalingam V, Girdhar A, Patel BK. Molecular Mechanisms of TDP-43 Misfolding and Pathology in Amyotrophic Lateral Sclerosis. Front Mol Neurosci 12, 25 (2019).

42. Santamaria N, Alhothali M, Alfonso MH, Breydo L, Uversky VN. Intrinsic disorder in proteins involved in amyotrophic lateral sclerosis. Cell Mol Life Sci 74, 1297–1318 (2017).

43. Zhang YJ, et al. Progranulin mediates caspase-dependent cleavage of TAR DNA binding protein-43. J Neurosci 27, 10530–10534 (2007).

44. Jiang LL, et al. Two mutations G335D and Q343R within the amyloidogenic core region of TDP-43 influence its aggregation and inclusion formation. Sci Rep 6, 23928 (2016).

45. Bolognesi B, Faure AJ, Seuma M, Schmiedel JM, Tartaglia GG, Lehner B. The mutational landscape of a prion-like domain. Nat Commun 10, 4162 (2019).

46. Lancaster AK, Nutter-Upham A, Lindquist S, King OD. PLAAC: a web and command-line application to identify proteins with prion-like amino acid composition. Bioinformatics 30, 2501–2502 (2014).

47. El-Gebali S, et al. The Pfam protein families database in 2019. Nucleic Acids Res 47, D427–D432 (2019).

48. Potter SC, Luciani A, Eddy SR, Park Y, Lopez R, Finn RD. HMMER web server: 2018 update. Nucleic Acids Res 46, W200–W204 (2018).

49. Hentze MW, Castello A, Schwarzl T, Preiss T. A brave new world of RNA-binding proteins. Nat Rev Mol Cell Biol 19, 327–341 (2018).

50. The UniProt C. UniProt: the universal protein knowledgebase. Nucleic Acids Res 45, D158–D169 (2017).

51. Dinkel H, et al. ELM 2016--data update and new functionality of the eukaryotic linear motif resource. Nucleic Acids Res 44, D294–300 (2016).

52. Yang X, et al. Stress granule-defective mutants deregulate stress responsive transcripts. PLoS Genet 10, e1004763 (2014).

53. Yang F, et al. Identifying pathogenicity of human variants via paralog-based yeast complementation. PLoS Genet 13, e1006779 (2017).

54. Steinmetz LM, et al. Systematic screen for human disease genes in yeast. Nat Genet 31, 400–404 (2002).

55. Couthouis J, et al. A yeast functional screen predicts new candidate ALS disease genes. Proc Natl Acad Sci U S A 108, 20881–20890 (2011).

56. Livi CM, Klus P, Delli Ponti R, Tartaglia GG. catRAPID signature: identification of ribonucleoproteins and RNA-binding regions. Bioinformatics 32, 773–775 (2016).

57. Buchan JR, Yoon JH, Parker R. Stress-specific composition, assembly and kinetics of stress granules in Saccharomyces cerevisiae. J Cell Sci 124, 228–239 (2011).

58. Li H, Shi, H., Zhu, Z., Wang, H., Niu, L., Teng, M. Crystal Structure of the First Two RRM Domains of Yeast Poly(U) Binding Protein (Pub1). RCSB PDB, (2010).

59. Sanchez de Groot N, Torrent Burgas M, Ravarani CN, Trusina A, Ventura S, Babu MM. The fitness cost and benefit of phase-separated protein deposits. Mol Syst Biol 15, e8075 (2019).

60. Gsponer J, Babu MM. Cellular strategies for regulating functional and nonfunctional protein aggregation. Cell reports 2, 1425–1437 (2012).

61. Wheeler JR, Matheny T, Jain S, Abrisch R, Parker R. Distinct stages in stress granule assembly and disassembly. Elife 5, (2016).

62. Klus P, Ponti RD, Livi CM, Tartaglia GG. Protein aggregation, structural disorder and RNA-binding ability: a new approach for physico-chemical and gene ontology classification of multiple datasets. BMC Genomics 16, 1071 (2015).

63. Tartaglia GG, Vendruscolo M. The Zyggregator method for predicting protein aggregation propensities. Chem Soc Rev 37, 1395–1401 (2008).

64. Wang M, Herrmann CJ, Simonovic M, Szklarczyk D, von Mering C. Version 4.0 of PaxDb: Protein abundance data, integrated across model organisms, tissues, and cell-lines. Proteomics 15, 3163–3168 (2015).

65. Oughtred R, et al. The BioGRID interaction database: 2019 update. Nucleic Acids Res 47, D529–D541 (2019).

66. Tartaglia GG, Pechmann S, Dobson CM, Vendruscolo M. Life on the edge: a link between gene expression levels and aggregation rates of human proteins. Trends Biochem Sci 32, 204–206 (2007).

67. Olzscha H, et al. Amyloid-like aggregates sequester numerous metastable proteins with essential cellular functions. Cell 144, 67–78 (2011).

68. Cerase A, Armaos A, Neumayer C, Avner P, Guttman M, Tartaglia GG. Phase separation drives X-chromosome inactivation: a hypothesis. Nat Struct Mol Biol 26, 331–334 (2019).

69. Mittag T, Parker R. Multiple Modes of Protein-Protein Interactions Promote RNP Granule Assembly. J Mol Biol 430, 4636–4649 (2018).

70. Maharana S, et al. RNA buffers the phase separation behavior of prion-like RNA binding proteins. Science 360, 918–921 (2018).

71. Garcia-Jove Navarro M, et al. RNA is a critical element for the sizing and the composition of phase-separated RNA-protein condensates. Nat Commun 10, 3230 (2019).

72. Hennig S, et al. Prion-like domains in RNA binding proteins are essential for building subnuclear paraspeckles. J Cell Biol 210, 529–539 (2015).

73. Tartaglia GG, Cavalli A, Pellarin R, Caflisch A. Prediction of aggregation rate and aggregation-prone segments in polypeptide sequences. Protein Sci 14, 2723–2734 (2005).

74. Cid F. Computational approaches to characterize RNP granules. Tesi Doctoral UPF, (2019).

75. Khong A, Matheny T, Jain S, Mitchell SF, Wheeler JR, Parker R. The Stress Granule Transcriptome Reveals Principles of mRNA Accumulation in Stress Granules. Mol Cell 68, 808–820 e805 (2017).

76. Saha S, et al. Polar Positioning of Phase-Separated Liquid Compartments in Cells Regulated by an mRNA Competition Mechanism. Cell 166, 1572–1584 e1516 (2016).

77. Falahati H, Pelham-Webb B, Blythe S, Wieschaus E. Nucleation by rRNA Dictates the Precision of Nucleolus Assembly. Curr Biol 26, 277–285 (2016).

78. Lee C, Occhipinti P, Gladfelter AS. PolyQ-dependent RNA-protein assemblies control symmetry breaking. J Cell Biol 208, 533–544 (2015).

79. Nishimoto Y, et al. The long non-coding RNA nuclear-enriched abundant transcript 1_2 induces paraspeckle formation in the motor neuron during the early phase of amyotrophic lateral sclerosis. Mol Brain 6, 31 (2013).

80. Lin Y, Schmidt BF, Bruchez MP, McManus CJ. Structural analyses of NEAT1 lncRNAs suggest long-range RNA interactions that may contribute to paraspeckle architecture. Nucleic Acids Res 46, 3742–3752 (2018).

81. Van Treeck B, Protter DSW, Matheny T, Khong A, Link CD, Parker R. RNA self-assembly contributes to stress granule formation and defining the stress granule transcriptome. Proc Natl Acad Sci U S A 115, 2734–2739 (2018).

82. Franzmann TM, Alberti S. Prion-like low-complexity sequences: Key regulators of protein solubility and phase behavior. J Biol Chem 294, 7128–7136 (2019).

83. Beckmann BM, et al. The RNA-binding proteomes from yeast to man harbour conserved enigmRBPs. Nat Commun 6, 10127 (2015).

84. Tartaglia GG, Caflisch A. Computational analysis of the S. cerevisiae proteome reveals the function and cellular localization of the least and most amyloidogenic proteins. Proteins 68, 273–278 (2007).

85. Brennan CM, et al. Protein aggregation mediates stoichiometry of protein complexes in aneuploid cells. Genes Dev 33, 1031–1047 (2019).

86. Qamar S, et al. FUS Phase Separation Is Modulated by a Molecular Chaperone and Methylation of Arginine Cation-pi Interactions. Cell 173, 720–734 e715 (2018).

87. An H, et al. ALS-linked FUS mutations confer loss and gain of function in the nucleus by promoting excessive formation of dysfunctional paraspeckles. Acta Neuropathol Commun 7, 7 (2019).

