## Supplementary Figures and Tables for "RNA-Binding and Prion Domains: The Yin and Yang of Phase Separation"

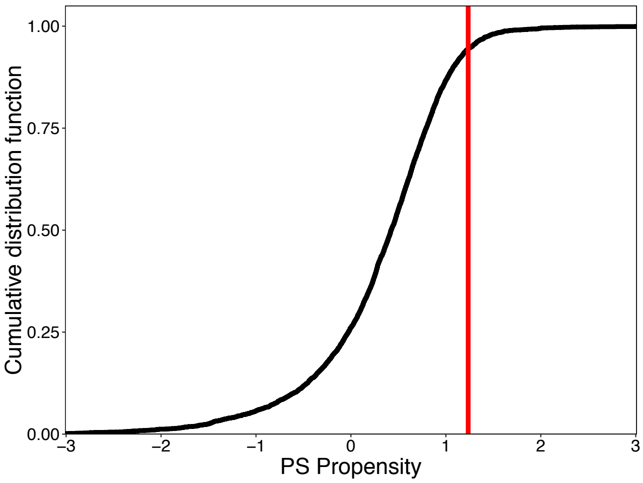


***Figure S1. Pub1 strong phase separation propensity among yeast proteome.*** *Cumulative distribution function (CDF) of the yeast proteome indicates that Pub1 has a high propensity to phase separate (red line).*

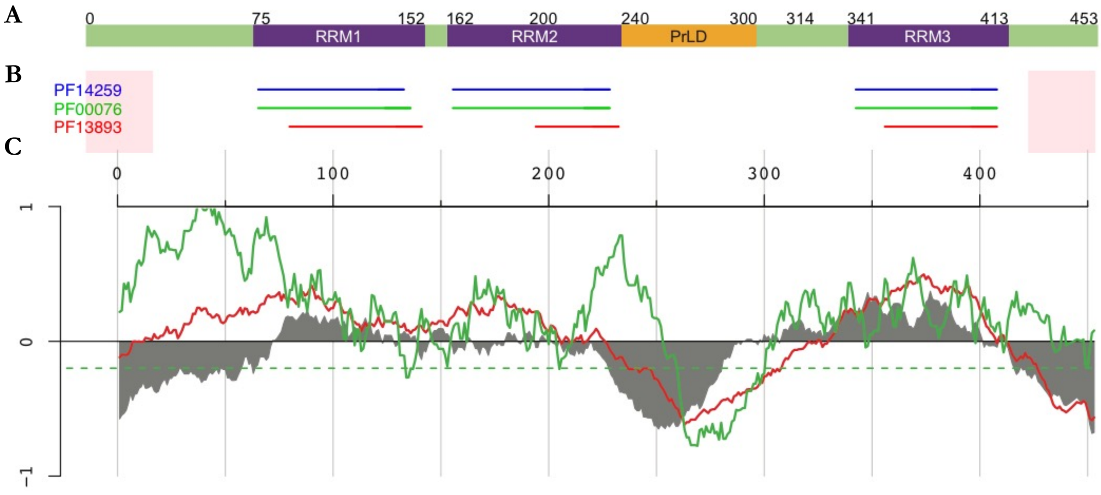


***Figure S2. Pub1 sequence analysis. A)*** *Pub1 sequence. RNA recognition motifs (RRM) in purple, PrLD in yellow.* ***B)*** *PFAM motifs highlighted with blue, green and red lines [1]. Pink areas indicate ends of the sequence.* ***C)*** *Predictions of prion-like regions of Pub1 with PLAAC [2] (red) and PAPA [3] (green) algorithms. Intrinsically unfolded regions are represented using negative values with FoldIndex tool [4] (grey).*

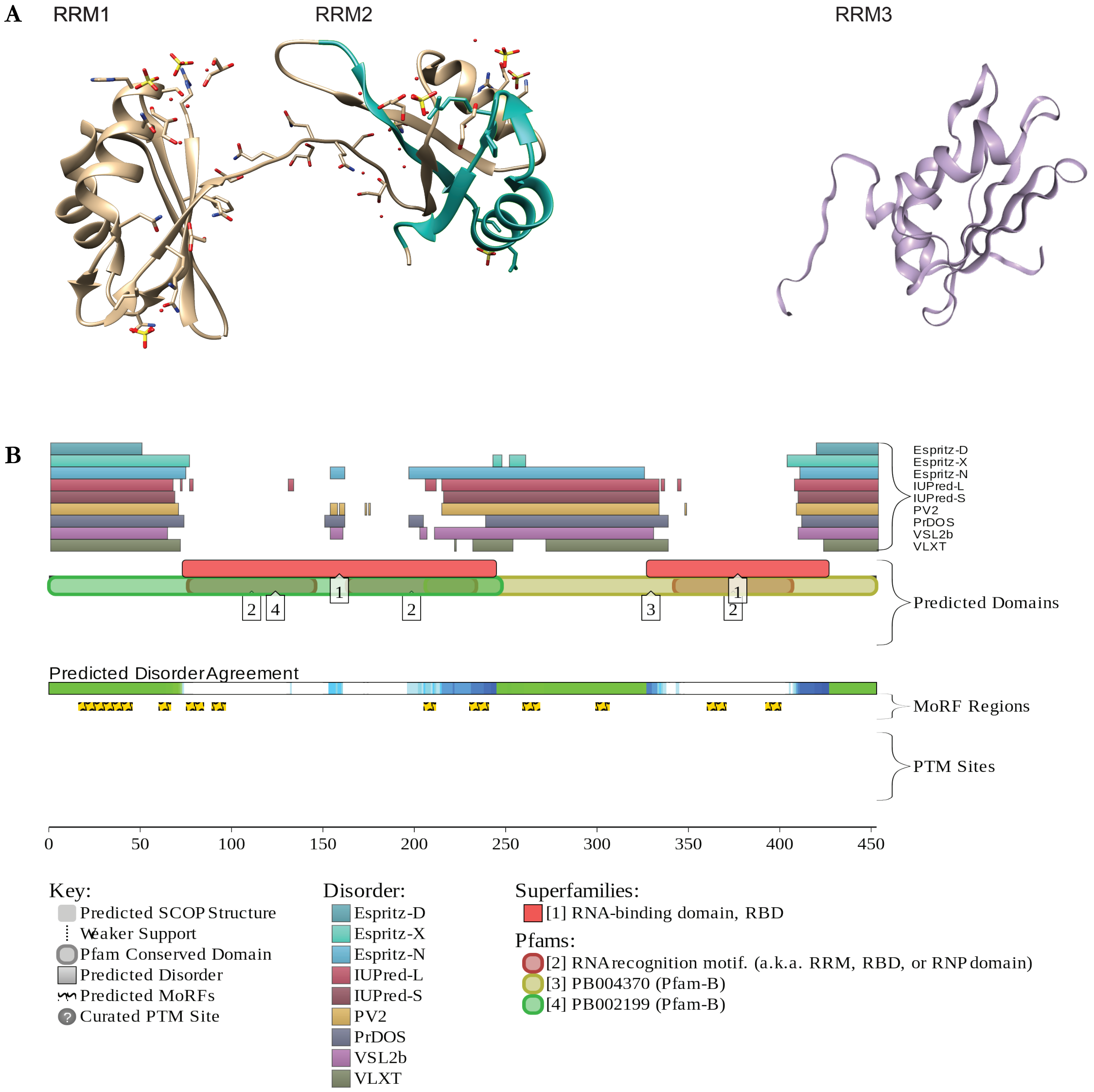


***Figure S3. Pub1 structured and disordered regions. A)** Crystal structures of RRM1-RRM2* [5] *(overlap with catGRANULE peak highlighted in blue) and RRM3* [6] *(PDB). **B)** Different predictions of disorder in Pub1 sequence, including a disorder agreement according to $D_2P_2$.*

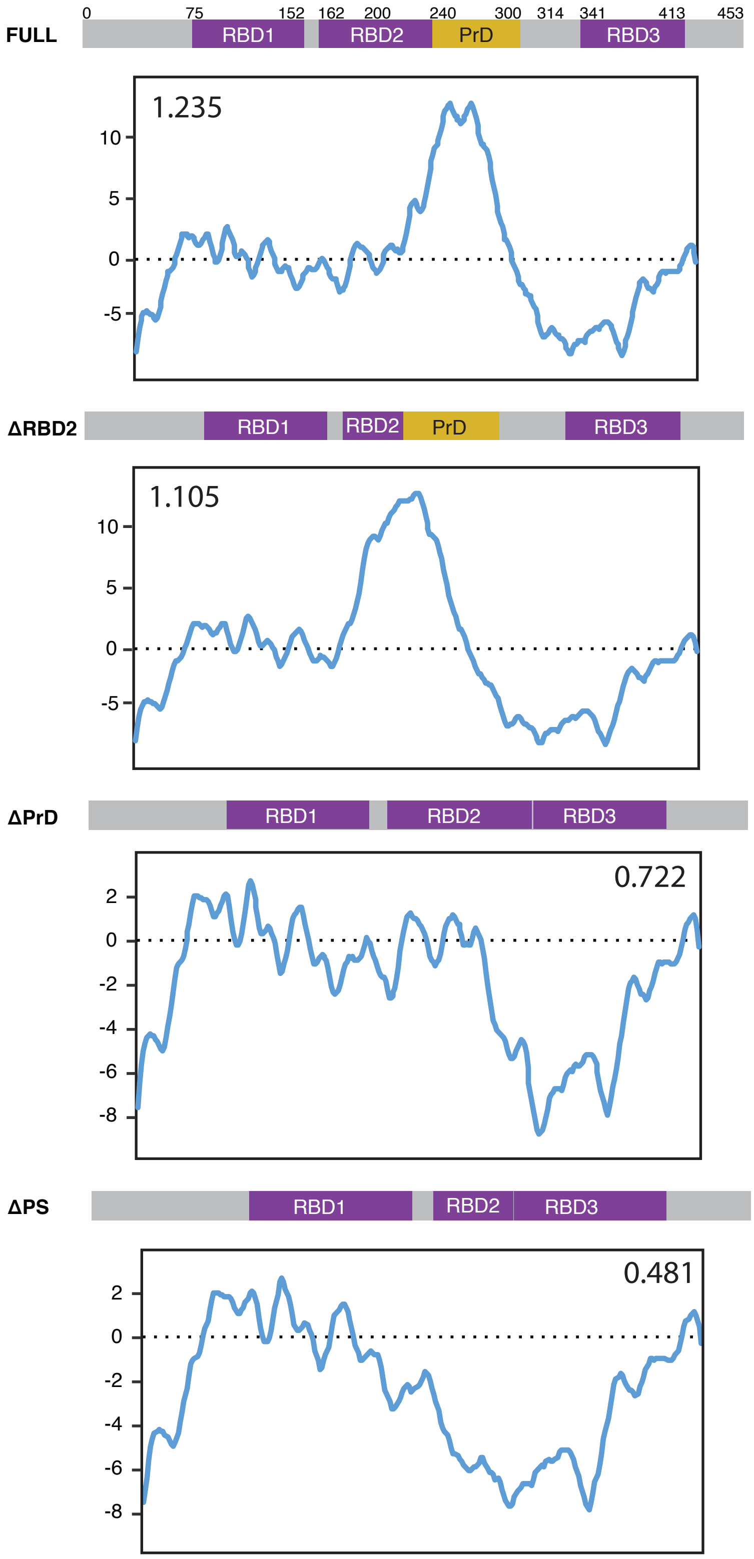


***Figure S4. PUB1 variants propensity to phase separate.*** *Score and profile of phase separation propensity predicted by catGRANULE along the four Pub1 variants analysed in the present work.*

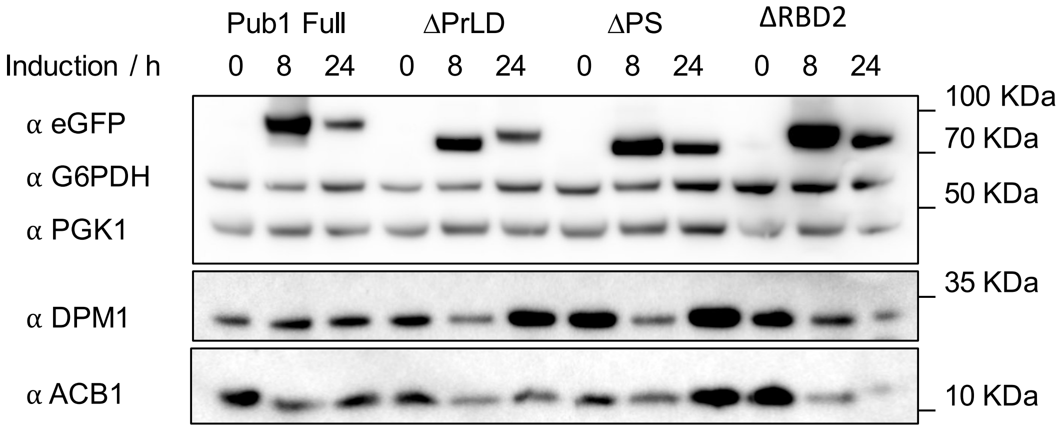


| | Pub1FULL | | ΔPrLD | | ΔPS | | ΔRBD2 | |
|---|---|---|---|---|---|---|---|---|
| **eGFP/G6PDH** | 8h | 24h | 8h | 24h | 8h | 24h | 8h | 24h |
| | 1.76 | 1.10 | 1.68 | 1.20 | 1.72 | 1.23 | 1.82 | 1.24 |

***Figure S5. Western blot indicates consistent expression between different Pub1 variants over time.*** *Up, antibody against eGFP showing the expression of Pub1-eGFP (78 KDa), ΔRRM2 (75 KDa), ΔPrLD-eGFP (70 KDa) and ΔPS-eGFP (67 KDa). Different housekeeping genes were evaluated to ensure a loading control no affected by the overexpression system. Down, quantification of the ratio between eGFP and G6PDH (eGFP/G6PDH).*

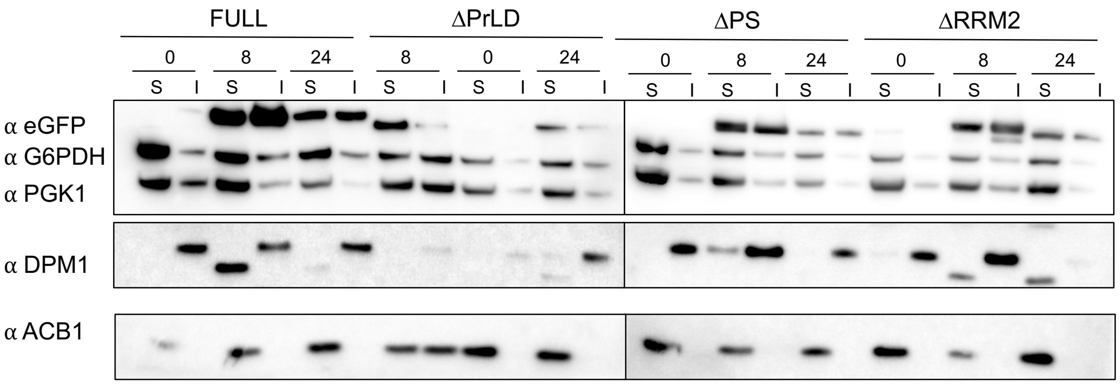


***Figure S6. Western blot indicates partition of Pub1 variants between soluble (S) and insoluble (I) fractions.*** *Antibody against eGFP show the expression of Pub1-eGFP (78 KDa), ΔRRM2 (75 KDa), ΔPrLD-eGFP (70 KDa) and ΔPS-eGFP (67 KDa). Different housekeeping genes were used to control for soluble and insoluble protein extractions.*

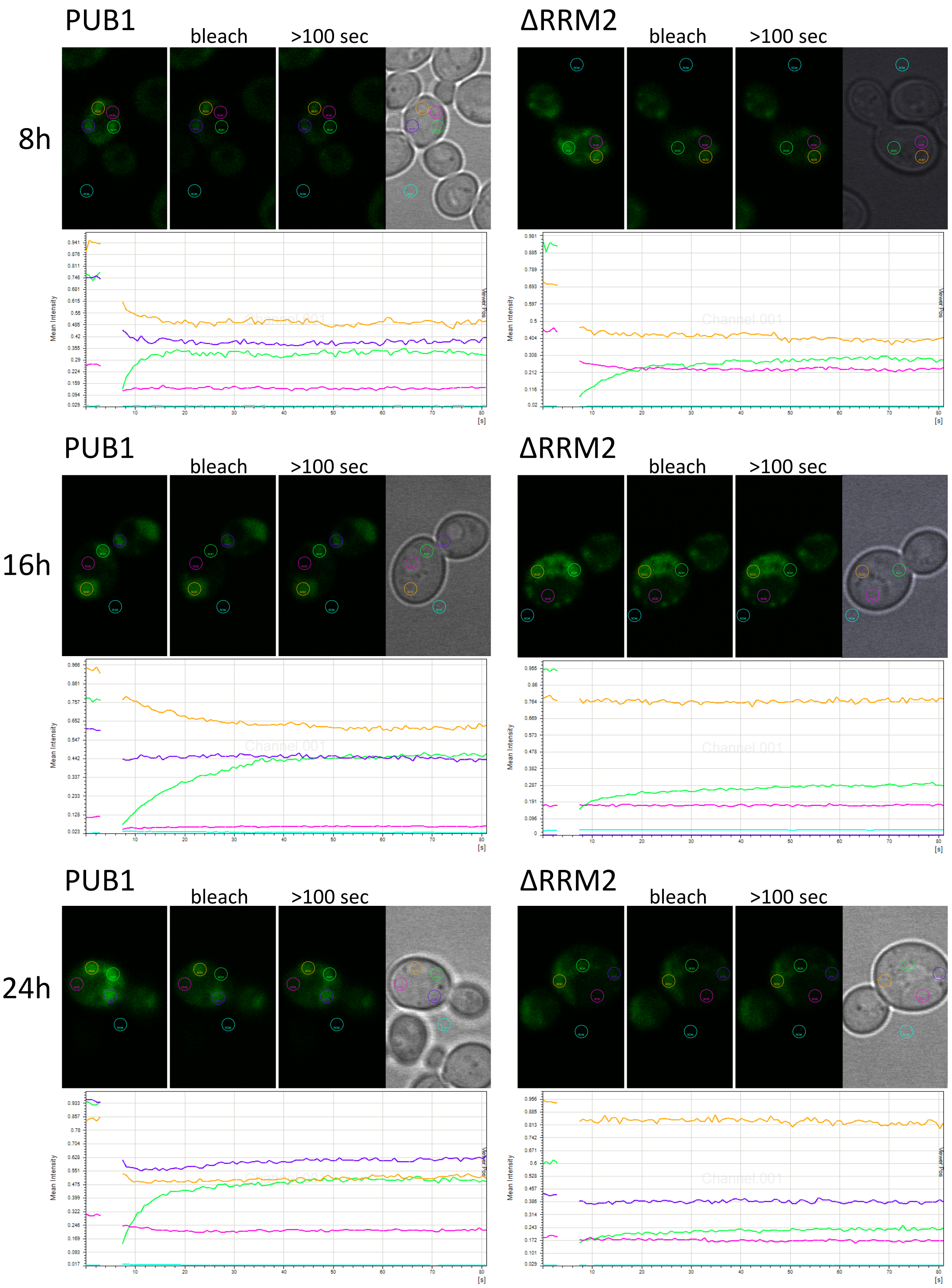


***Figure S7. FRAP experiments.*** *Confocal microscope images from left to right: immediately before bleach, immediately after bleach, 100 seconds after bleach and bright field. Green: bleached area; pink: cytoplasm background; yellow: not-bleached condensate; purple: not-bleached condensate; cyan: background outside cells. Plots indicate normalized fluorescence of each area (arbitrary units) over time (seconds). One example of each strain and condition is shown as example.*

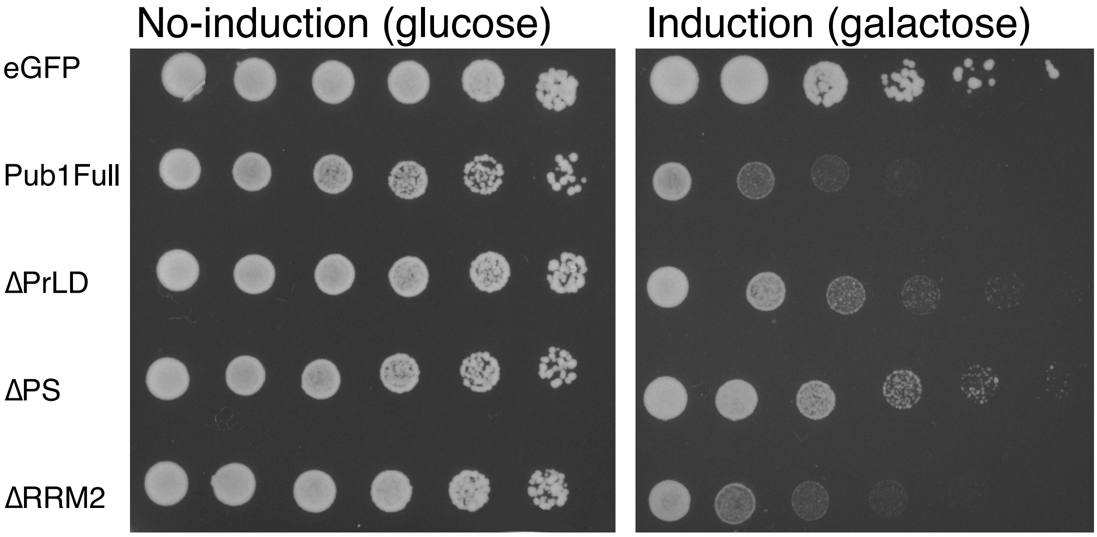


**Figure S8. Spot dilution reveals fitness impairment due to overexpression of Pub1 variants.** Cells were pre-incubated overnight in no-induction conditions and serially diluted from OD 0.1 (first drop) to 0.0016 (last drop) prior to agar plating. Image taken after three days of incubation at 30ºC.

| | | | LAG TIME (h) | | DOUBLING TIME (h) | | SATURATION OD | |
|---|---|---|---|---|---|---|---|---|
| | | | MEAN | SD | MEAN | SD | MEAN | SD |
| TOXICITY | 0h | FULL | 22.7677 | 3.1247 | 5.0688 | 1.0754 | 0.0638 | 0.0031 |
| | | DRRM2 | 20.6802 | 2.5188 | 3.8916 | 0.1393 | 0.1255 | 0.0454 |
| | | DPrLD | 22.2943 | 0.9217 | 3.8531 | 0.0673 | 0.1514 | 0.0232 |
| | | DPS | 19.3404 | 2.2043 | 3.7227 | 0.4654 | 0.3299 | 0.0709 |
| | | BKG | 14.9513 | 2.4142 | 2.0566 | 0.1553 | 0.8265 | 0.0112 |
| | 8h | FULL | 13.6464 | 1.0611 | 5.6197 | 1.4009 | 0.0906 | 0.0537 |
| | | DRRM2 | 13.2003 | 0.9704 | 4.1031 | 0.4596 | 0.1560 | 0.0214 |
| | | DPrLD | 11.0237 | 1.0526 | 3.7603 | 0.6485 | 0.3785 | 0.0412 |
| | | DPS | 11.1816 | 0.7206 | 3.6347 | 0.6079 | 0.4816 | 0.0360 |
| | | BKG | 6.9586 | 0.2282 | 1.9902 | 0.1172 | 0.8230 | 0.0279 |
| | 16h | FULL | 16.8348 | 1.5021 | 6.1272 | 1.2919 | 0.0211 | 0.0032 |
| | | DRRM2 | 14.0725 | 1.7179 | 4.3291 | 0.6793 | 0.0541 | 0.0252 |
| | | DPrLD | 13.3222 | 1.8738 | 3.8758 | 0.5408 | 0.1447 | 0.0766 |
| | | DPS | 12.3544 | 1.6242 | 3.1944 | 0.1607 | 0.2778 | 0.0720 |
| | | BKG | 8.5583 | 1.1406 | 1.9736 | 0.0544 | 0.8803 | 0.0090 |
| | 24h | FULL | NA | NA | 7.2612 | 0.6950 | 0.0133 | 0.0008 |
| | | DRRM2 | 13.8370 | 0.3517 | 5.4867 | 1.2897 | 0.0338 | 0.0096 |
| | | DPrLD | 15.9454 | 1.1422 | 4.9436 | 1.9845 | 0.0562 | 0.0038 |
| | | DPS | 11.2305 | 1.9466 | 3.4848 | 0.5227 | 0.3706 | 0.0355 |
| | | BKG | 5.0825 | 0.6495 | 1.8440 | 0.3437 | 0.8436 | 0.0407 |

**Table S1. Quantification of three parameters from Pub1 growth curves upon protein induction: lag time, doubling time and saturation peak.** Four Pub1 variants and background were assessed after different pre-induction times. All times are indicated in hours. Mean corresponds to arithmetic average of three independent biological replicates and corresponding standard deviations (SD) are calculated.

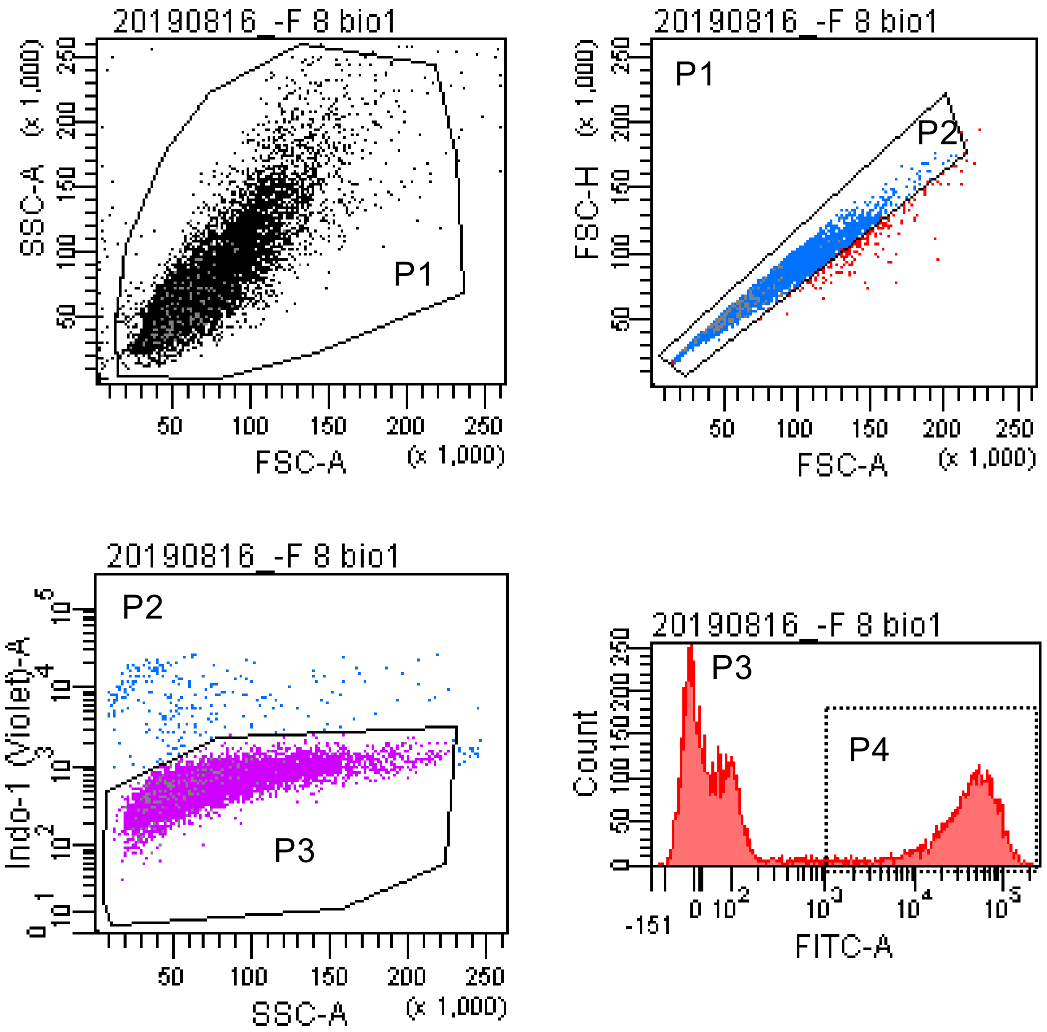


**Figure S9. Flow cytometer analysis indicates the percentage of cells expressing Pub1 variants fused to eGFP.** The figure depicts an example of gating process of Pub1 full strain after 8 hours of induction. P1: exclusion of debris. P2: exclusion of doublets from P1. P3: exclusion of dead cells from P2; corresponds to live cells population. P4: green fluorescent cells from P3; corresponds to cells expressing Pub1 variants.

| | | | % Dead cells | |
|---|---|---|---|---|
| | | | MEAN | SD |
| MORTALITY | 0 h | Full | 5.03% | 0.0126 |
| | | ΔRRM2 | 4.90% | 0.0105 |
| | | ΔPrD | 4.88% | 0.0096 |
| | | ΔPS | 5.42% | 0.0129 |
| | 8 h | Full | 2.77% | 0.0108 |
| | | ΔRRM2 | 3.29% | 0.0188 |
| | | ΔPrD | 2.78% | 0.0129 |
| | | ΔPS | 3.21% | 0.0112 |
| | 16 h | Full | 4.62% | 0.0021 |
| | | ΔRRM2 | 4.83% | 0.0080 |
| | | ΔPrD | 4.03% | 0.0120 |
| | | ΔPS | 3.81% | 0.0167 |
| | 24 h | Full | 4.00% | 0.0178 |
| | | ΔRRM2 | 4.41% | 0.0139 |
| | | ΔPrD | 4.88% | 0.0156 |
| | | ΔPS | 4.07% | 0.0349 |

| | | | % eGFP cells | |
|---|---|---|---|---|
| | | | MEAN | SD |
| EXPRESSION | 0 h | Full | 0.10% | 0.4443 |
| | | ΔRRM2 | 0.23% | 0.2082 |
| | | ΔPrD | 0.20% | 0.4970 |
| | | ΔPS | 0.23% | 0.2081 |
| | 8 h | Full | 39.13% | 4.6715 |
| | | ΔRRM2 | 42.07% | 1.7780 |
| | | ΔPrD | 46.43% | 5.7387 |
| | | ΔPS | 52.11% | 1.3350 |
| | 16 h | Full | 20.10% | 7.3920 |
| | | ΔRRM2 | 19.47% | 4.5010 |
| | | ΔPrD | 28.01% | 2.5848 |
| | | ΔPS | 35.72% | 2.2978 |
| | 24 h | Full | 8.17% | 4.9756 |
| | | ΔRRM2 | 6.50% | 4.3643 |
| | | ΔPrD | 8.48% | 4.0709 |
| | | ΔPS | 4.55% | 7.0377 |

**Table S2. Cell counting with flow cytometry.** Left, percentage of dead cells for each Pub1 variant overtime. The measured fraction corresponds to [(P2-P3)/P2] in Figure s9. Right, percentage of fluorescent cells for each Pub1 variant over time. Corresponds to [P4/P3] accordingly to Figure S9. Mean of 3 biological replicates. SD: standard deviation.

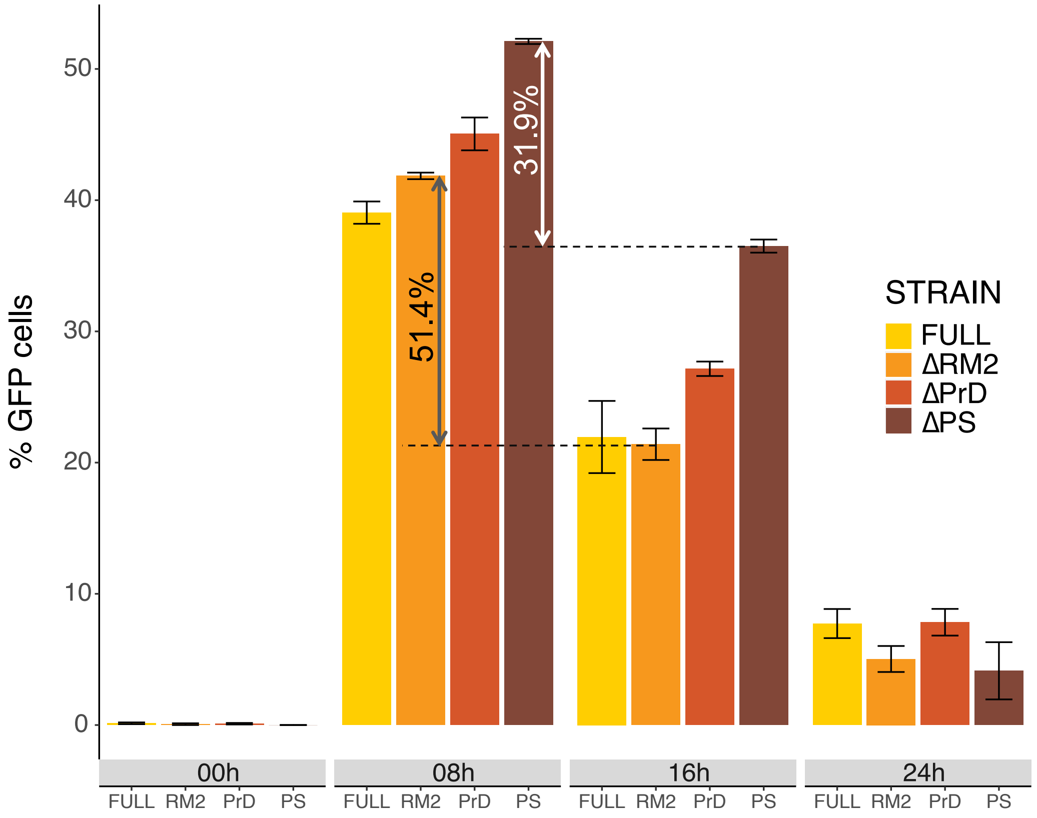


**Figure S9. Analysis of percentage of green fluorescence as a measure of Pub1 variant expression and plasmid retention.** Total percentage of fluorescent cells for each strain grouped by induction times. Quantifications are performed on selected gate P4 (green fluorescent live cells) for all conditions.

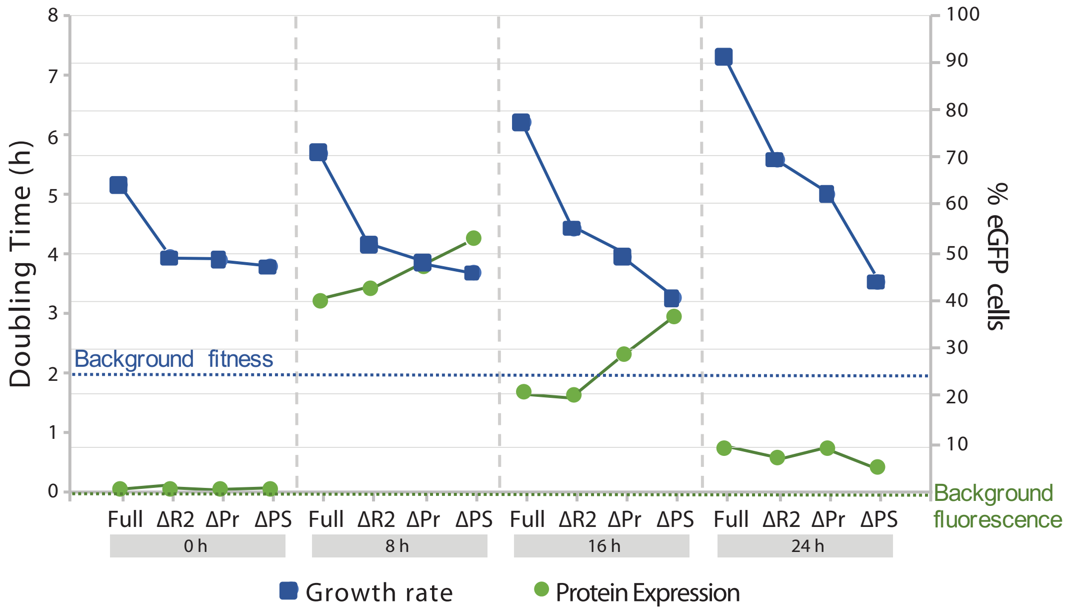


**Figure S11. Comparison of Pub1 variants expression and strain fitness.** Green: percentage of fluorescent cells in a population. Blue: time required (hours) for a strain to double its population. Dotted lines indicate background. The deceleration of cellular division cannot be explained just by plasmid loss (inability to produce uracil). The differences in plasmid retention between strains (35.72% ΔPS and 20.10% Pub1 full, after 16h of induction) do not matches with the dramatic differences in fitness (3.48 h DT ΔPS and 7.26 h DT full, after 24 h of induction).

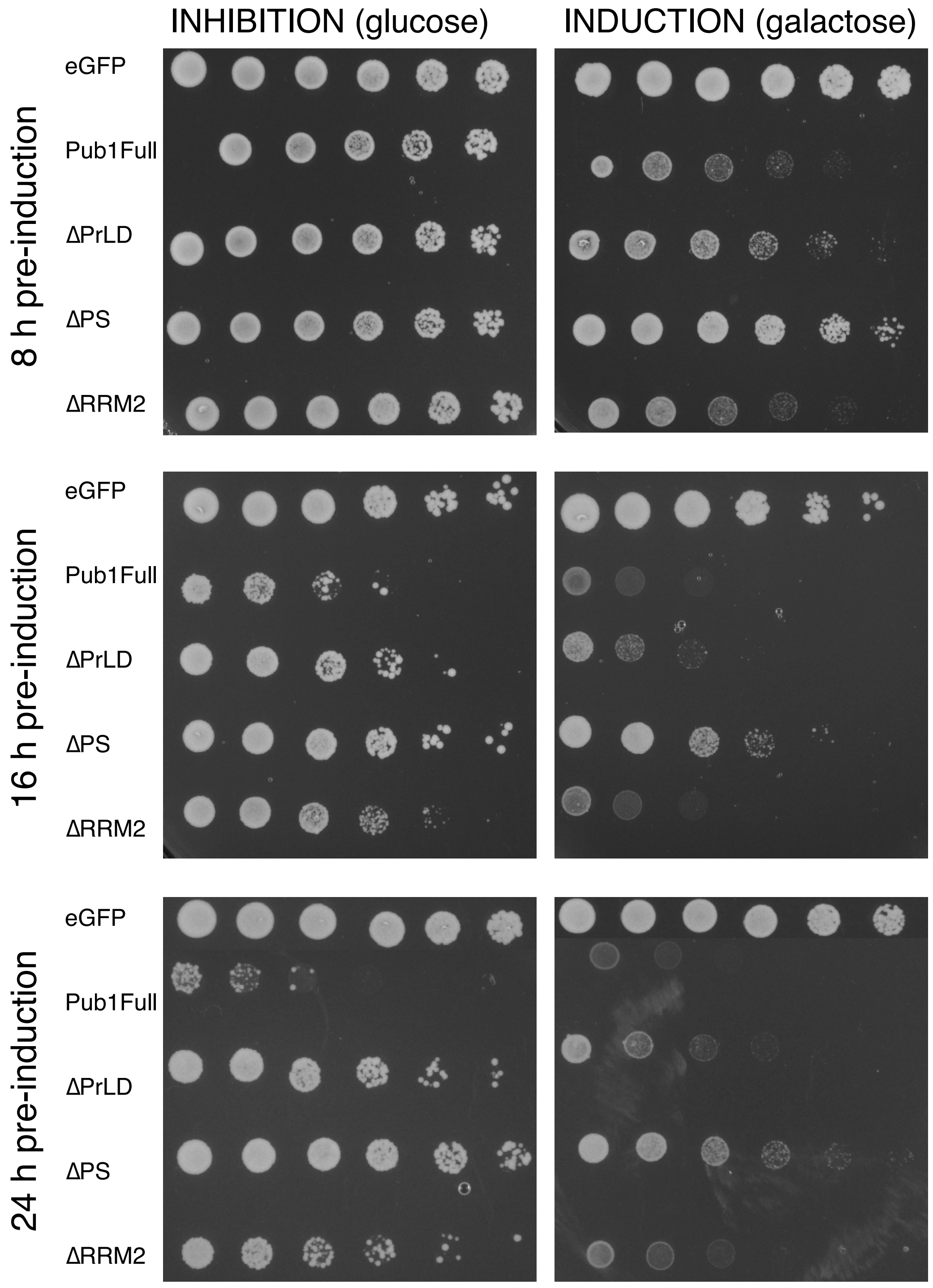


**Figure S12. Spot dilution reveals total recovery of Pub1 variants fitness after 8 h of pre-induction, partial recovery after 16 h and worse recovery after 24 h of pre-induction.** Cells were serially diluted from OD 0.1 (first drop) to 0.0016 (last drop) prior to agar plating. Images taken after three days of incubation at 30ºC.

| | | | LAG TIME (h) | | DOUBLING TIME (h) | | SATURATION OD | |
|---|---|---|---|---|---|---|---|---|
| | | | MEAN | SD | MEAN | SD | MEAN | SD |
| RECOVERY | 0h | FULL | 7.1781 | 1.3092 | 1.8542 | 0.0827 | 0.6550 | 0.0490 |
| | | DRR2 | 6.9217 | 1.4859 | 1.9534 | 0.0574 | 0.6178 | 0.0248 |
| | | DPrD | 8.1111 | 0.6797 | 1.6951 | 0.0220 | 0.6397 | 0.0381 |
| | | DPS | 6.1565 | 0.8581 | 1.9264 | 0.0710 | 0.6303 | 0.0304 |
| | | BKG | 7.0358 | 2.3304 | 1.5913 | 0.1142 | 0.6653 | 0.0550 |
| | 8h | FULL | 8.6897 | 0.9316 | 2.1158 | 0.0775 | 0.6172 | 0.0191 |
| | | DRR2 | 8.0158 | 0.8980 | 2.2580 | 0.1487 | 0.5821 | 0.0293 |
| | | DPrD | 6.6478 | 0.8185 | 2.1325 | 0.0518 | 0.6283 | 0.0218 |
| | | DPS | 6.2186 | 1.1683 | 2.0989 | 0.0417 | 0.6139 | 0.0193 |
| | | BKG | 5.3493 | 0.4695 | 1.4999 | 0.0512 | 0.7087 | 0.0243 |
| | 16h | FULL | 16.5501 | 1.3763 | 3.1324 | 0.3019 | 0.5872 | 0.0341 |
| | | DRR2 | 11.2851 | 2.9556 | 4.2024 | 0.1757 | 0.5013 | 0.0488 |
| | | DPrD | 9.2787 | 1.8371 | 2.8721 | 0.6240 | 0.6365 | 0.0183 |
| | | DPS | 8.4711 | 2.7200 | 2.6031 | 0.1943 | 0.6253 | 0.0226 |
| | | BKG | 6.5448 | 0.4896 | 1.4994 | 0.0598 | 0.7655 | 0.0269 |
| | 24h | FULL | 23,9448 | 6.3896 | 2.7259 | 0.1935 | 0.5278 | 0.0594 |
| | | DRR2 | 18.3000 | 1.6016 | 4.3054 | 0.5265 | 0.4484 | 0.0527 |
| | | DPrD | 13.6050 | 0.8328 | 2.8152 | 0.6948 | 0.5882 | 0.0353 |
| | | DPS | 9.7338 | 1.9226 | 2.8023 | 0.1507 | 0.5644 | 0.0292 |
| | | BKG | 4.3902 | 0.3715 | 1.4947 | 0.0786 | 0.7614 | 0.0319 |

**Table S3. Quantification of three parameters from Pub1 growth curves upon protein inhibition: lag time, doubling time and saturation peak.** Four Pub1 variants and background were assessed after different pre-induction times. All times are indicated in hours. Mean corresponds to arithmetic average of three independent biological replicates and corresponding standard deviations (SD) are calculated.

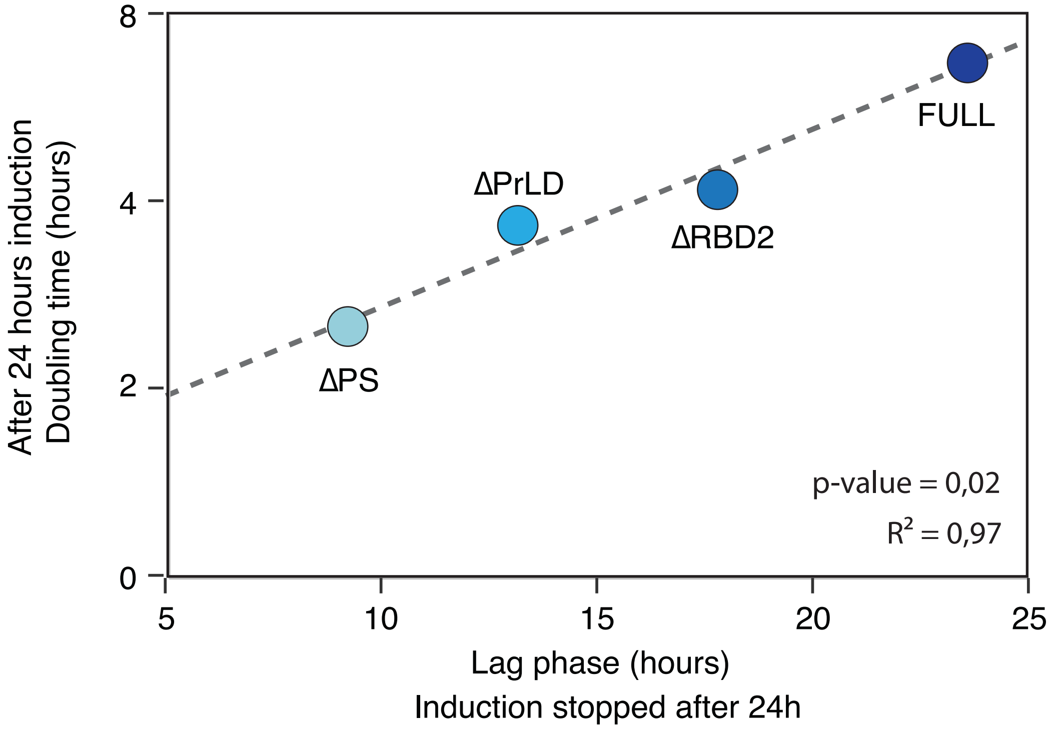


**Figure S13. Growth impairment correlates with the time to achieve the top division speed after stop induction.** Linear correlation between the doubling time at induction conditions and the lag time measured after stop the induction. In both cases the pre-growth was 24 hours in presence of galactose as inductor.

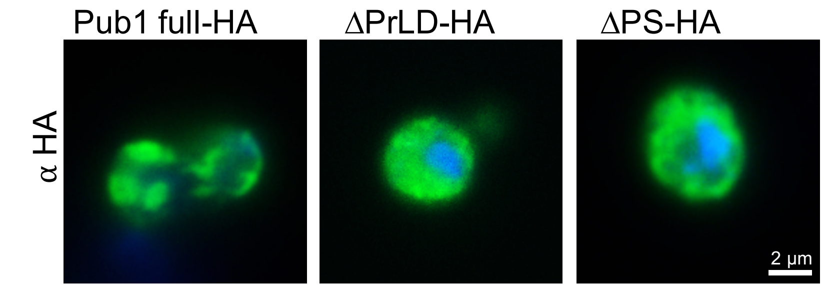


**Figure S14. Immunofluorescence images indicate that Pub1 Full-HA tagged forms condensates.** ΔPrLD-HA and ΔPS-HA present a diffused pattern. In green protein variant tagged (HA antibody); in blue nuclei stained (DAPI).

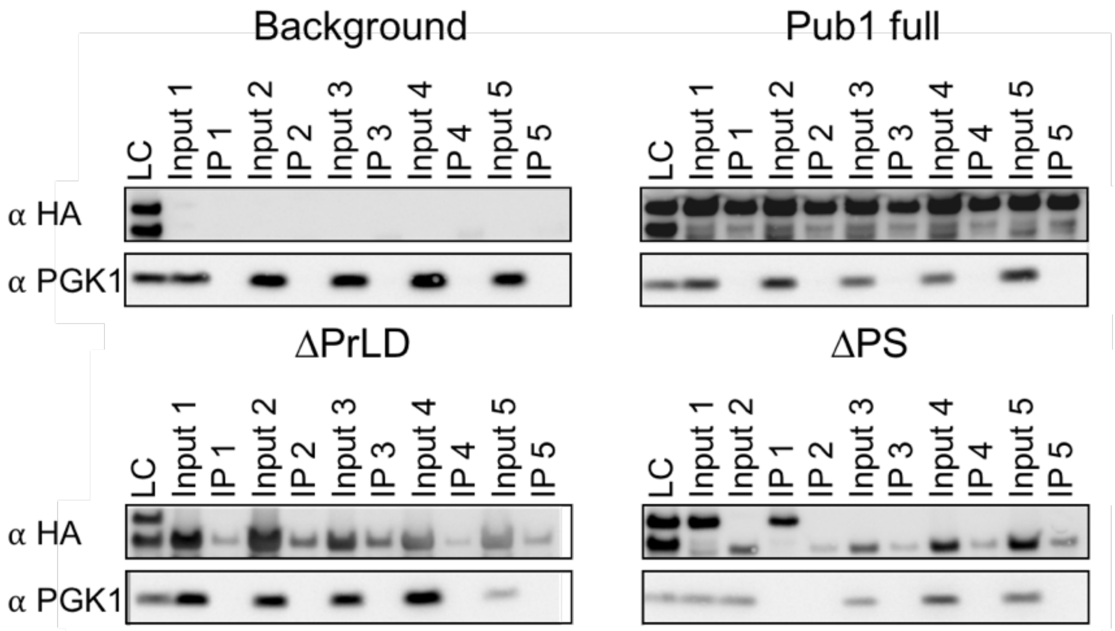


**Figure S15. Western blots confirm the quality of the immunoprecipitations of Pub1 variants.** Five immunoprecipitations of each strain were analysed with antibodies against HA tag and Pgk1 housekeeping protein. Protein samples before (input) and after (IP) immunoprecipitation revealed sufficient Pub1 variants extraction to proceed with mass spectrometry. A loading control (LC, mix of equal parts of four strains input1) was loaded in all membranes in order to compare them. Identification of incorrect protein size of ΔPS sample 1 allowed to discard it for further analysis.

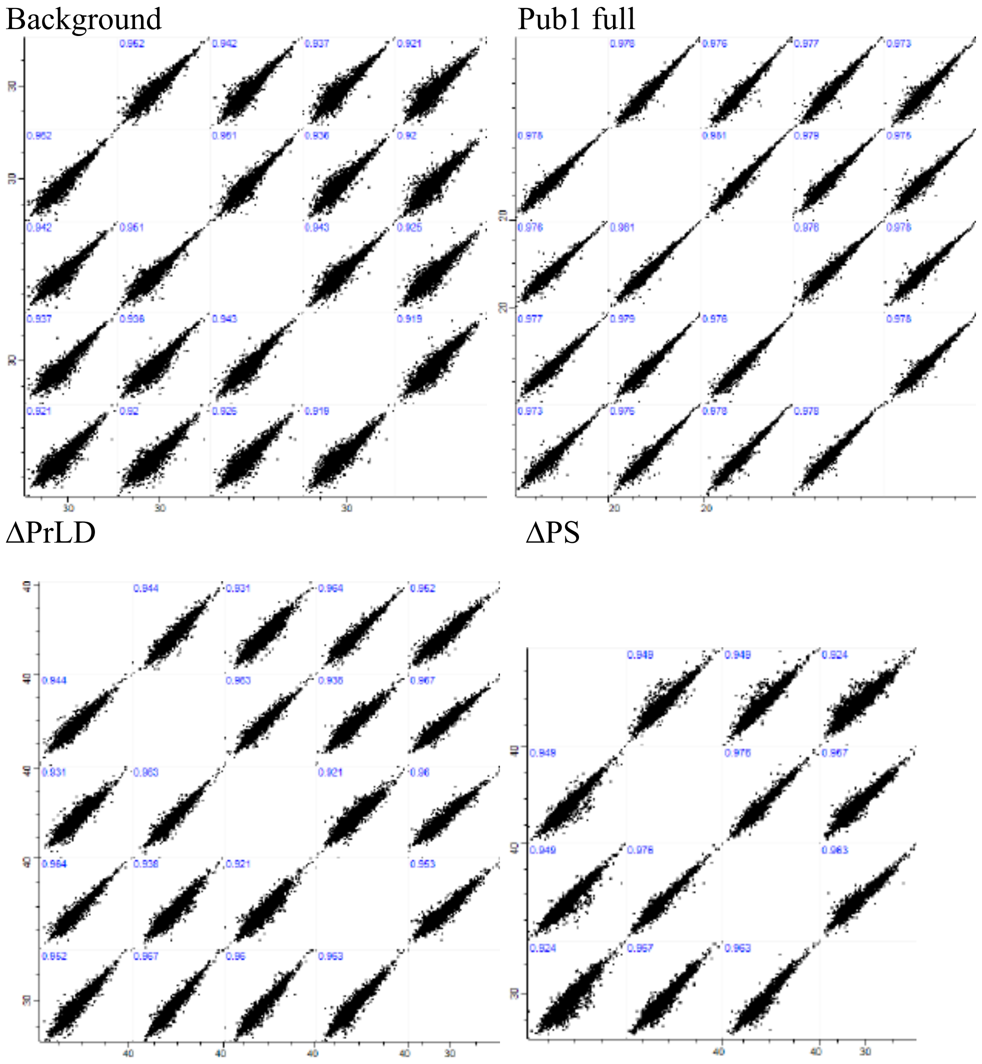


**Figure S16. Mass spectrometry results indicate good correlation between all biological replicates of Pub1 variants.** Pearson correlation (log2 LFQ (label free quantifications)) between different biological replicates indicated in blue.

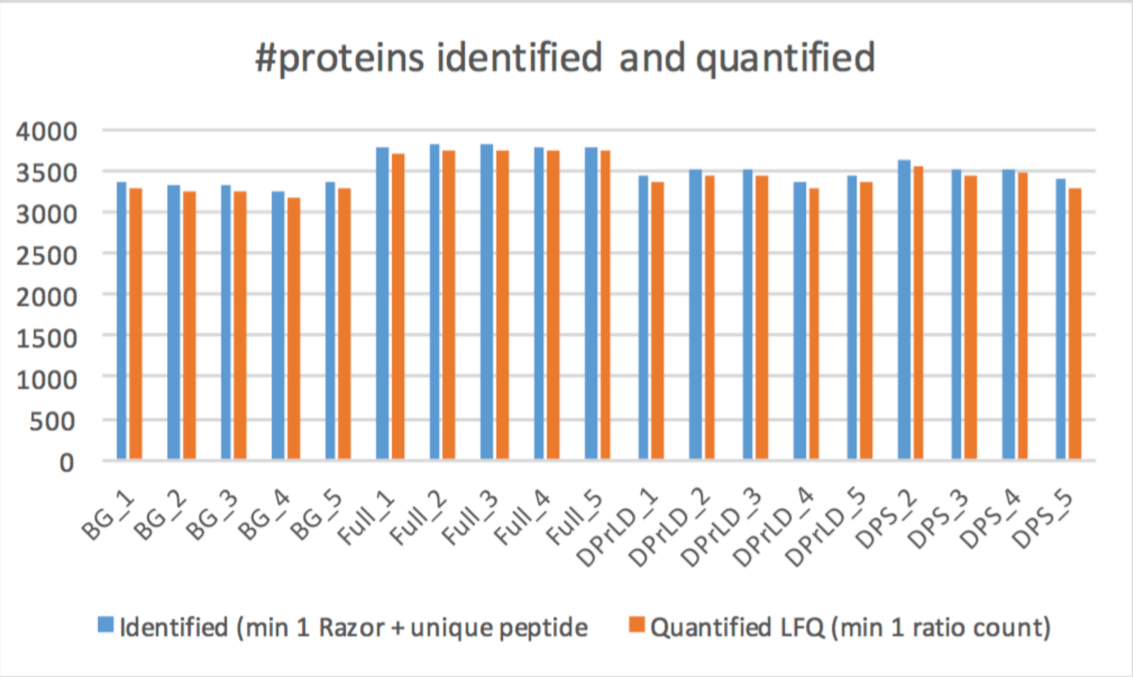


**Figure S17. Mass spectrometry detection of immunoprecipitated proteins.** The number of identified and quantified proteins is similar in all immunoprecipitated samples. SAMPLES: BG = background, Full = Pub1full, ΔPrLD, ΔPS

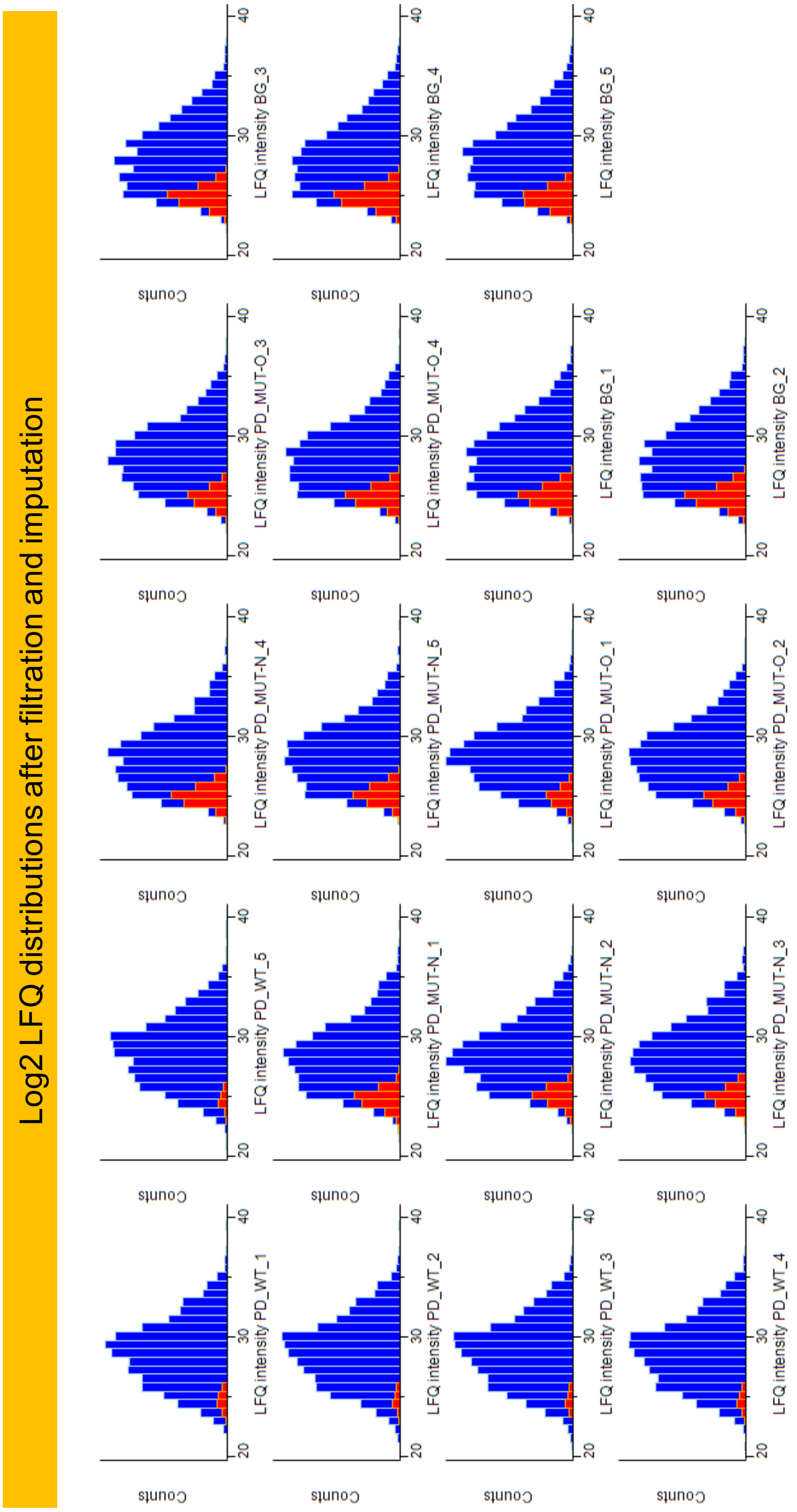


**Figure S18. Bioinformatic data analysis on mass spectrometry results.** The data was filtered for proteins having at least 3 valid LFQ values in at least one sample group (red). Analysis using Perseus (version 1.5.2.6). WT: Pub1 full; MUT-N: ΔPrLD; MUT-O: ΔPS.

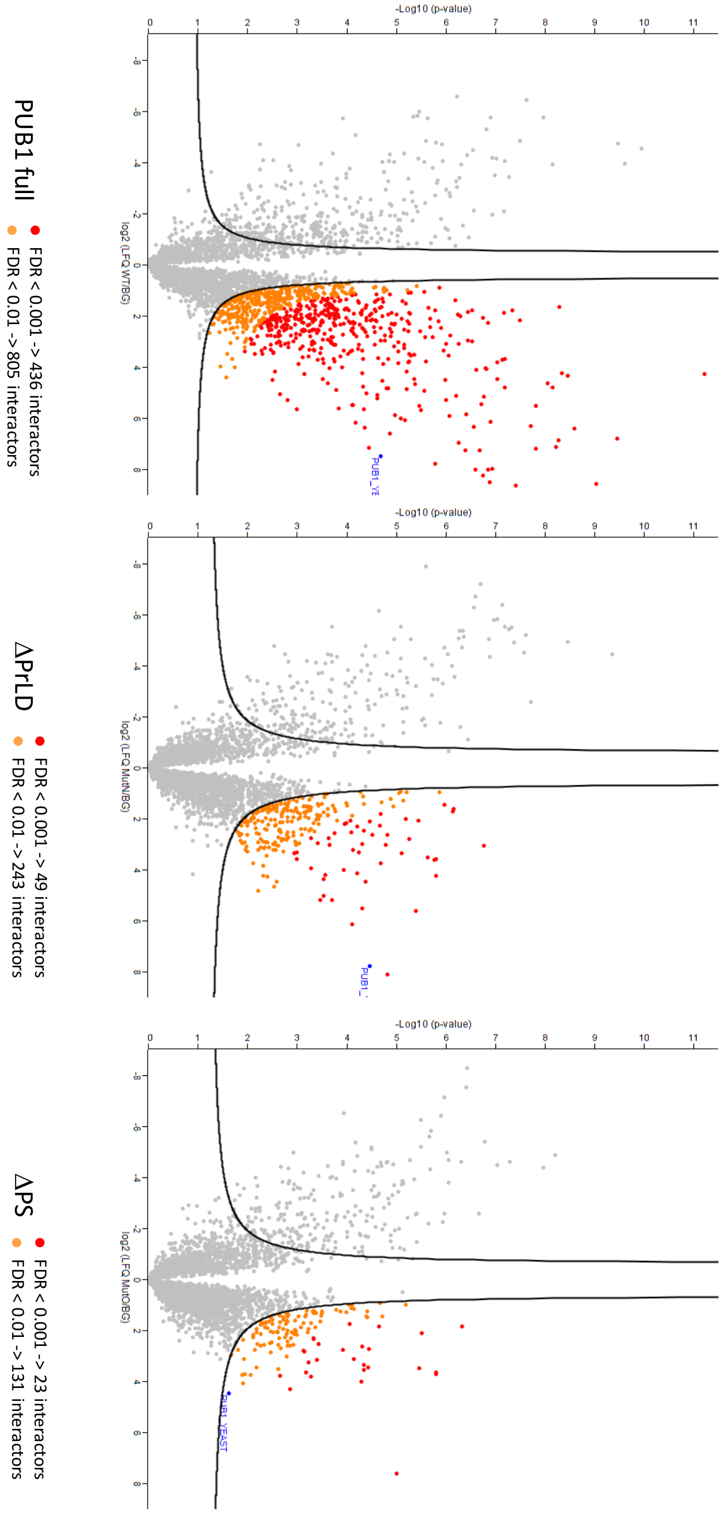


**Figure S19. Volcano plots represent identified interactors of Pub1 variants.** P-value versus LFQ ratio (sample/background), in logarithmic scales. Statistical analysis: 2-samples t-test, FDR threshold at 0.01 (orange) or 0.001 (red).

**TABLE S4. Mass spectrometry results indicate the identity of interactors of Pub1 variants.** Proteins with False Discovery Range (FDR) <0.001 are highlighted in red. BG corresponds to background (no Pub1, no HA tag).

See file Table.S4.xlsx

**Table S5. List of proteins with RBDs and PRDs.**

| id | Total prediction Score | LLR | catGranule | GranulePrD | hmm_present |
|---|---|---|---|---|---|
| sp.P02381.RMAR_YEAST | 0.73 | 35.863 | 1.85398 | no overlap | 0 |
| sp.P05453.ERF3_YEAST | 0.52 | 42.019 | 1.71388 | overlap | 2 |
| sp.P06634.DED1_YEAST | 0.74 | 25.177 | 1.97798 | overlap | 0 |
| sp.P09547.SWI1_YEAST | 0.45 | 42.003 | 1.27658 | overlap | 0 |
| sp.P11746.MCM1_YEAST | 0.72 | 43.643 | 0.504423 | overlap | 0 |
| sp.P11792.SCH9_YEAST | 0.59 | 30.277 | 1.35063 | NoPRD | 0 |
| sp.P12383.PDR1_YEAST | 0.29 | 28.03 | 0.875973 | no overlap | 0 |
| sp.P14680.YAK1_YEAST | 0.58 | 28.611 | 0.869177 | no overlap | 0 |
| sp.P14922.CYC8_YEAST | 0.65 | 52.523 | 0.0366241 | overlap | 0 |
| sp.P18480.SNF5_YEAST | 0.63 | 62.818 | 0.605474 | no overlap | 0 |
| sp.P18494.GLN3_YEAST | 0.71 | 26.504 | 1.16467 | no overlap | 0 |
| sp.P18899.DDR48_YEAST | 0.33 | 25.173 | 4.77004 | overlap | 0 |
| sp.P19659.MED15_YEAST | 0.7 | 44.643 | 0.817868 | overlap | 0 |
| sp.P22082.SNF2_YEAST | 0.67 | 28.447 | 1.26481 | no overlap | 0 |
| sp.P23202.URE2_YEAST | 0.51 | 28.319 | 0.808932 | overlap | 0 |
| sp.P23250.RPI1_YEAST | 0.59 | 31.451 | 0.837993 | NoPRD | 0 |
| sp.P23291.KC11_YEAST | 0.55 | 27.195 | 1.0745 | no overlap | 0 |
| sp.P23292.KC12_YEAST | 0.67 | 29.726 | 1.39578 | no overlap | 0 |
| sp.P23900.FPS1_YEAST | 0.45 | 27.485 | 1.06435 | NoPRD | 0 |
| sp.P24276.SSD1_YEAST | 0.56 | 25.578 | 1.22093 | no overlap | 1 |
| sp.P25299.RNA15_YEAST | 0.32 | 32.565 | 0.762262 | NoPRD | 1 |
| sp.P25339.PUF4_YEAST | 0.52 | 30.93 | 0.937476 | no overlap | 8 |
| sp.P25367.RNQ1_YEAST | 0.82 | 46.717 | 3.11099 | overlap | 0 |
| sp.P27895.CIN8_YEAST | 0.34 | 25.653 | 1.15965 | no overlap | 0 |
| sp.P29295.HRR25_YEAST | 0.44 | 28.227 | 0.817165 | no overlap | 0 |
| sp.P31384.CCR4_YEAST | 0.4 | 32.388 | 0.877198 | overlap | 0 |
| sp.P32505.NAB2_YEAST | 0.85 | 49.903 | 0.498264 | overlap | 0 |
| sp.P32521.PAN1_YEAST | 0.65 | 44.576 | 1.32066 | overlap | 0 |
| sp.P32588.PUB1_YEAST | 0.81 | 40.622 | 1.23508 | overlap | 3 |
| sp.P32770.NRP1_YEAST | 0.6 | 48.64 | 1.6159 | overlap | 0 |
| sp.P32896.PDC2_YEAST | 0.39 | 27.69 | 0.812771 | overlap | 0 |
| sp.P33417.IXR1_YEAST | 0.84 | 40.682 | 0.71533 | overlap | 0 |
| sp.P34217.PIN4_YEAST | 0.61 | 25.937 | 0.905148 | no overlap | 1 |
| sp.P34756.FAB1_YEAST | 0.54 | 33.406 | 1.28805 | no overlap | 0 |
| sp.P34758.SCD5_YEAST | 0.55 | 26.646 | 0.866482 | overlap | 0 |
| sp.P34761.WHI3_YEAST | 0.68 | 25.654 | 0.950593 | no overlap | 1 |

| | | | | | |
|---|---|---|---|---|---|
| sp.P35732.DEF1_YEAST | 0.88 | 50.369 | 0.970604 | overlap | 0 |
| sp.P38042.CDC27_YEAST | 0.62 | 32.425 | 0.501286 | overlap | 0 |
| sp.P38080.AKL1_YEAST | 0.68 | 34.195 | 1.27204 | no overlap | 0 |
| sp.P38114.TBS1_YEAST | 0.58 | 26.265 | 0.748197 | NoPRD | 0 |
| sp.P38129.TAF5_YEAST | 0.51 | 31.64 | 1.32833 | no overlap | 6 |
| sp.P38180.YBI1_YEAST | 0.56 | 37.441 | 0.652357 | overlap | 0 |
| sp.P38216.YBM6_YEAST | 0.67 | 50.371 | 1.06685 | overlap | 0 |
| sp.P38266.AIM3_YEAST | 0.59 | 30.521 | 1.376 | no overlap | 0 |
| sp.P38330.RMD9L_YEAST | 0.47 | 36.04 | 0.572787 | overlap | 0 |
| sp.P38429.SAP30_YEAST | 0.45 | 42.301 | 1.90574 | overlap | 0 |
| sp.P38691.KSP1_YEAST | 0.63 | 31.877 | 1.73236 | no overlap | 0 |
| sp.P38856.AP18A_YEAST | 0.6 | 30.468 | 0.674246 | overlap | 0 |
| sp.P38996.NAB3_YEAST | 0.89 | 32.37 | 1.30876 | no overlap | 1 |
| sp.P39008.POP2_YEAST | 0.48 | 29.83 | 0.499865 | no overlap | 1 |
| sp.P39081.PCF11_YEAST | 0.56 | 26.439 | 0.635482 | no overlap | 0 |
| sp.P39523.YM11_YEAST | 0.7 | 29.791 | 1.41359 | no overlap | 0 |
| sp.P40002.MIT1_YEAST | 0.77 | 51.243 | 1.31636 | overlap | 0 |
| sp.P40070.LSM4_YEAST | 0.64 | 34.622 | 2.01803 | overlap | 1 |
| sp.P40356.MED3_YEAST | 0.79 | 34.367 | 0.379342 | overlap | 0 |
| sp.P40467.ASG1_YEAST | 0.48 | 57.091 | 0.984424 | overlap | 0 |
| sp.P40485.SLM1_YEAST | 0.46 | 31.912 | 0.589155 | overlap | 0 |
| sp.P40956.GTS1_YEAST | 0.67 | 32.081 | 0.908644 | no overlap | 0 |
| sp.P43582.WWM1_YEAST | 0.59 | 27.211 | 1.59994 | overlap | 0 |
| sp.P45978.SCD6_YEAST | 0.88 | 25.072 | 1.52838 | overlap | 0 |
| sp.P47135.JSN1_YEAST | 0.55 | 26.776 | 0.899529 | no overlap | 4 |
| sp.P48562.CLA4_YEAST | 0.37 | 28.409 | 0.677399 | no overlap | 0 |
| sp.P50109.PSP2_YEAST | 0.88 | 33.891 | 2.7844 | overlap | 0 |
| sp.P50896.PSP1_YEAST | 0.34 | 26.769 | 1.39897 | no overlap | 0 |
| sp.P53165.SGF73_YEAST | 0.82 | 26.427 | 1.6984 | overlap | 0 |
| sp.P53829.CAF40_YEAST | 0.32 | 26.623 | 0.481602 | overlap | 0 |
| sp.P53894.CBK1_YEAST | 0.45 | 49.359 | 0.918935 | overlap | 0 |
| sp.P54785.MOT3_YEAST | 0.74 | 33.517 | 1.26381 | no overlap | 0 |
| sp.Q02629.NU100_YEAST | 0.32 | 29.606 | 2.99684 | overlap | 0 |
| sp.Q02630.NU116_YEAST | 0.55 | 36.748 | 3.15392 | overlap | 0 |
| sp.Q02792.XRN2_YEAST | 0.62 | 31.857 | 1.1286 | overlap | 1 |
| sp.Q03735.NAB6_YEAST | 0.55 | 27.156 | 1.43868 | no overlap | 1 |
| sp.Q03761.TAF12_YEAST | 0.5 | 39.389 | 0.774268 | no overlap | 0 |
| sp.Q03825.MSS11_YEAST | 0.83 | 56.531 | 1.18398 | no overlap | 0 |
| sp.Q04195.SIZ1_YEAST | 0.79 | 29.749 | 0.861647 | no overlap | 1 |
| sp.Q04978.YMF3_YEAST | 0.62 | 27.853 | 0.937264 | NoPRD | 0 |
| sp.Q05785.ENT2_YEAST | 0.67 | 46.562 | 0.552057 | overlap | 0 |
| sp.Q05854.YL278_YEAST | 0.6 | 41.417 | 1.0284 | overlap | 0 |

| | | | | | |
|---|---|---|---|---|---|
| sp.Q06449.PIN3_YEAST | 0.57 | 33.407 | 0.537094 | overlap | 0 |
| sp.Q06628.ATG13_YEAST | 0.52 | 29.869 | 1.04539 | no overlap | 0 |
| sp.Q07807.PUF3_YEAST | 0.47 | 27.552 | 1.40874 | no overlap | 6 |
| sp.Q08601.MCA1_YEAST | 0.47 | 34.95 | 1.27127 | overlap | 0 |
| sp.Q08732.HRK1_YEAST | 0.73 | 35.348 | 0.911045 | NoPRD | 0 |
| sp.Q08925.MRN1_YEAST | 0.55 | 32.593 | 1.09863 | no overlap | 4 |
| sp.Q08954.YP199_YEAST | 0.61 | 26.537 | 1.11904 | NoPRD | 0 |
| sp.Q08972.NEW1_YEAST | 0.43 | 40.246 | 1.23107 | no overlap | 0 |
| sp.Q12124.MED2_YEAST | 0.56 | 37.309 | 2.133 | overlap | 0 |
| sp.Q12139.YP022_YEAST | 0.37 | 29.607 | 0.939273 | overlap | 0 |
| sp.Q12151.UPC2_YEAST | 0.39 | 26.606 | 0.924146 | no overlap | 0 |
| sp.Q12221.PUF2_YEAST | 0.53 | 54.609 | 1.03568 | overlap | 4 |
| sp.Q12224.RLM1_YEAST | 0.8 | 42.846 | 1.69378 | no overlap | 0 |
| sp.Q12361.GPR1_YEAST | 0.71 | 55.706 | 1.27369 | no overlap | 0 |
| sp.Q12518.ENT1_YEAST | 0.62 | 25.069 | 0.992314 | no overlap | 0 |
| sp.Q99383.HRP1_YEAST | 0.84 | 30.267 | 2.23889 | overlap | 2 |

Pub1 Full

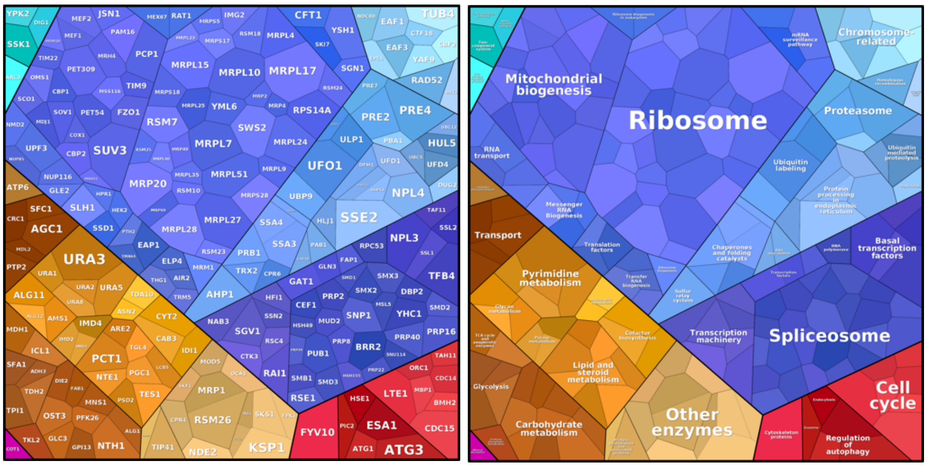


ΔPrLD

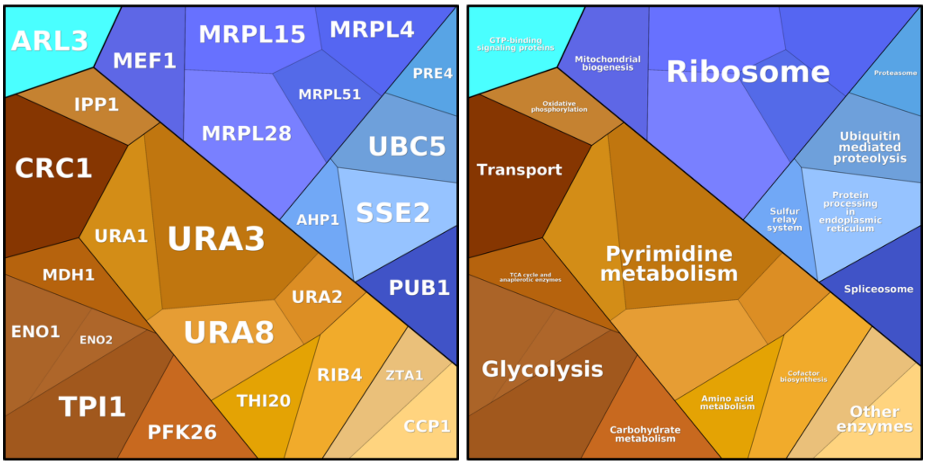


ΔPS

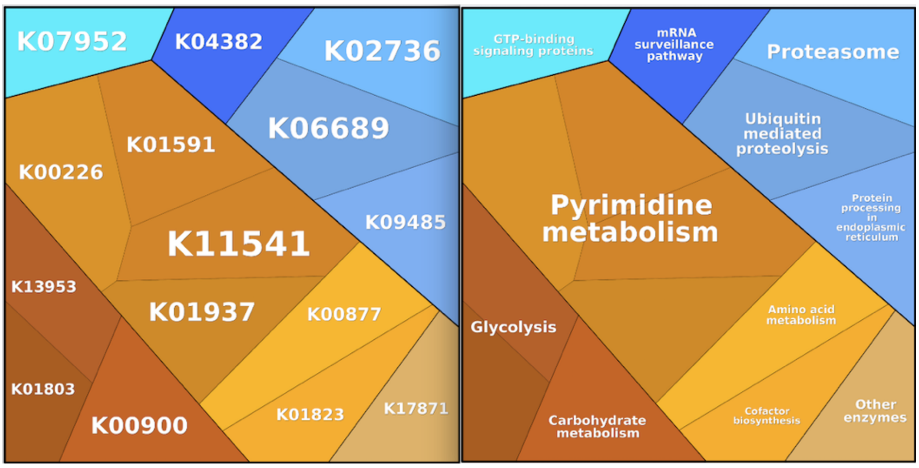


**Figure S19. Interaction networks of Pub1 variants visualized with Proteomaps.** Proteomaps show the quantitative composition of proteomes with a focus on protein function. They are built automatically from proteome data and based on the KEGG Pathways gene classification. Each protein is shown by a polygon and functionally related proteins are arranged in common regions. To emphasize their interactivity with PUB1, polygon areas represent the different amount between the sample and the background measured by mass spectrometry (p-values from Table S4).

**Table S7. Annotation of proteins found in the mass-spec experiments.** Name (Uniprot), type of strain (1=Full, 2=ΔPrLD, 3= ΔPS), essential or not (1/0), length, PS score, number of physical interactors, prion propensity, abundance and reported in SG or not (1/0).

See file Table.S7.xlsx

1. Livi CM, Klus P, Delli Ponti R, Tartaglia GG. catRAPID signature: identification of ribonucleoproteins and RNA-binding regions. *Bioinformatics* **32**, 773-775 (2016).

2. Lancaster AK, Nutter-Upham A, Lindquist S, King OD. PLAAC: a web and command-line application to identify proteins with prion-like amino acid composition. *Bioinformatics* **30**, 2501-2502 (2014).

3. Toombs JA, Petri M, Paul KR, Kan GY, Ben-Hur A, Ross ED. De novo design of synthetic prion domains. *Proc Natl Acad Sci U S A* **109**, 6519-6524 (2012).

4. Prilusky J*, et al.* FoldIndex: a simple tool to predict whether a given protein sequence is intrinsically unfolded. *Bioinformatics* **21**, 3435-3438 (2005).

5. Li H, Shi, H., Zhu, Z., Wang, H., Niu, L., Teng, M. Crystal Structure of the First Two RRM Domains of Yeast Poly(U) Binding Protein (Pub1). *RCSB PDB*, (2010).

6. Santiveri CM, Mirassou Y, Rico-Lastres P, Martinez-Lumbreras S, Perez-Canadillas JM. Pub1p C-terminal RRM domain interacts with Tif4631p through a conserved region neighbouring the Pab1p binding site. *PLoS One* **6**, e24481 (2011).
