## Supplementary material for "RNA-Binding and Prion Domains: The Yin and Yang of Phase Separation": materials and methods

*Material and methods*

### Datasets

#### Proteomes

Human and Yeast proteomes were obtained from UniProt [1]. Sequence files containing all canonical sequences from each organism's reference proteome were obtained from the UniProt website. This resulted in 20,404 canonical and reviewed human proteins (proteome UP000005640 from UniProt) and 6,049 canonical and reviewed yeast proteins (proteome UP000002311 from UniProt release 2018/11, strain ATCC 204508 / S288c).

Annotations of experimentally validated yeast stress granule proteins were obtained from Jain *et al.*, 2016 [2]. Annotations of experimentally validated prion proteins were obtained from Alberti *et al.*, 2009 [3].

A confident list of yeast essential list was used according to Pache *et al.*, 2009 [4] (combining data from Steinmetz *et al.*, 2002 [5] and Ghaemmaghami *et al.,* 2003 [6]). Dosage sensitive proteins dataset resulted from the combination of Sopko *et al.*, 2006 [7] and Makanae *et al.*, 2013 [8].

A very recent census of proteins with experimental evidence of RNA-binding activity in human (1914 known RBPs) and yeast (1273 known RBPs) was used to flag proteins as known RBPs (from Hentze *et al.*, 2018 [9]. In addition, location of RBDs (Pfam [10] annotated) within RBPs was screened using HMMER [11].

#### *cat*GRANULE analysis

*cat*GRANULE was employed to identify proteins assembling into biological condensates. Scores > 0 indicate that a protein is prone to phase separate. Structural disorder, nucleic acid binding propensity and amino acid patterns such as arginine-glycine and phenylalanine-glycine are key features combined in this computational approach [12].

The protein part considered prone to phase separate and used in the overlap analysis (**Figure 1**), is defined as the region of positive values around the highest peak of the *cat*GRANULE profile. For proteins lacking positive scores, the region is defined as the 50 amino acids centred around the highest score.

#### PLAAC analysis

Prion-Like Amino Acid Composition (PLAAC) from Lancaster *et al.* 2014 [13] searches protein sequences to identify probable prion sub-sequences using a hidden-Markov model (HMM) algorithm. The algorithm was downloaded from https://github.com/whitehead/plaac and run locally to our servers for the Human and

Yeast proteome. Default settings except for window w=50 (to match *cat*GRANULE window) were used. Considered PrLD regions for following analysis were defined as [PRDstart, PRDend]. Proteins not predicted to contain a PrLD are assigned as "No PrLD".

**Overlap analysis**

Phase separation regions (highest *cat*GRANULE [12] peak) were systematically screened for the overlap with annotated RBDs (Pfamv [10]). Similarly, phase separation regions were screened for the overlap with PrLDs (PLAAC [13]). Two regions were considered as overlapping when at least 20% of the shortest region length coincides with the other region.

**Yeast strains and genetic engineering**

S. cerevisiae strain S288c BY4741 (MATa his3Δ1 leu2Δ0 met15Δ0 ura3Δ0) was used for all experiments. Uracil auxotrophy was used as selection marker for all genetic engineering (including the gene URA along the cloning sequence).

The Gal-1 promoters were amplified via tool-box PCR as designed by Janke *et al.*, 2004 [14]. Amplified fragments were genome integrated via a standard lithium acetate transformation protocol and the correct insertion of the promoter was verified by PCR on extracted genomic DNA.

For plasmid overexpression experiments, a strain lacking the PUB1 gene was used (ATCC, from yeast deletion project). Pub1 full was expressed via cloning into p416-Gal1 and p426-Gal1 (Addgene). eGFP alone was expressed into p426-Gal1. For the characterization of Pub1 deletion variants, p416-Gal1-PUB1full-eGFP was linearized and reannealed (Gibson reaction) to delete corresponding sequence fragments. Oligonucleotides (primers) used for cloning are listed below; forward (F) and reverse (R)are written from 5' to 3'; in grey: flanking regions for recombination; lower case: spacer before tag (three alanines) or stop codon after tag.

Primers to linearize p416 or p426:
F: GCAACAGCAGCAACAACAAgccgcagcgATGGTGAGCAAGGGCGAGGAGCT
R: TCTTCGTTATTTTCAGACATCTCCTTGACGTTAAAGTATAGAGGT

Primers to amplify PUB1 full gene from BY4741:
F: TATACTTTAACGTCAAGGAGATGTCTGAAAATAACGAAGAACAAC
R: CCTCGCCCTTGCTCACCATcgctgcggcTTGTTGTTGCTGCTGTTGCTGCTG

Primers to linearize p416-PUB1full for ΔRRM2:
F: ATTATTGTTATTGTTATCACGACCAGTTTGCATGTCCCACATAACATGACC

R: CGTGATAACAATAACAATAATAATTATCAACAGCGTCGTAACTACGG

Primers to linearize p416-PUB1full for ΔPrLD:
F: AACTGGGCTGCTAAGCAATCCCAGACCATTGGTTTACCTC
R: CAATGGTCTGGGATTGCTTAGCAGCCCAGTTGATTCTTAG

Primers to linearize p416-PUB1full for ΔPS:
F: AGACTTCCCAACCCCAACAATCCCAGACCATTGGTTTACC
R: TTGTTGGGGTTGGGAAGTCTTTGAAGGCATTCCTTAAAGT

For localization and immunochemistry analysis, the genes were tagged with either eGFP, mCherry or hemagglutinin (HA) by means of p416 plasmid cloning. For co-localization studies, strains from the GFP collection[15] were used. Primers used for cloning are listed below (same format code as previously used).

Primers to linearize p416-PUB1full deleting eGFP tag (HA swap):
F: TATGACGTTCCAGATTACGCtagTAATTAGTTATGTCACGCTTAC
R: TCGTATGGGTAACCCATTCCcgctgcggcTTGTTGTTGCTGCTGT

Primers to amplify HA gene from template:
F: AGCAACAACAAgccgcagcgGGAATGGGTTACCCATACGATGTTC
R: CGTGACATAACTAATTActaAGCGTAATCTGGAACGTCATATGGA

Primers to linearize p416-PUB1full deleting eGFP tag (mCherry swap):
F: TGGACGAGCTGTACAAGtagTCATGTAATTAGTTATGTCACGCTTAC
R: TCCTCGCCCTTGCTCACcatCGCTGCGGCTTGTTGTTG

Primers to amplify mCherry gene from template:
F: ATGGTGAGCAAGGGCGAGGAGGATA
R: CTACTTGTACAGCTCGTCCATGCCG

PUB1 Variants

**Red** RRM2
**Green** Prion until the end of PS pic
**Bold** Remaining 14 aa of the prion

>FULL
MSENNEEQHQQQQQQQPVAVETPSAVEAPASADPSSEQSVAVEGNSEQAEDN
QGENDPSV

VPANAITGGRETSDRVLYVGNLDKAITEDILKQYFQVGGPIANIKIMIDKNNKN
VNYAFV
EYHQSHDANIALQTLNGKQIENNIVKINWAFQSQQSSSDDTFNLFVGDLNVNV
DDETLRN
AFKDFPSYLSGHVMWDMQTGSSRGYGFVSFTSQDDAQNAMDSMQGQDLNGR
PLRINWAAK
RDNNNNNNNYQQRRNYGNNNRGGFRQYNSNNNNNMNMGMNMNMNMNMNN
SRGMPPSSMGMP
**IGAMPLPSQGQPQQ**SQTIGLPPQVNPQAVDHIIRSAPPRVTTAYIGNIPHFATEA
DLIPL
FQNFGFILDFKHYPEKGCCFIKYDTHEQAAVCIVALANFPFQGRNLRTGWGKER
SNFMPQ
QQQQGGQPLIMNDQQQPVMSEQQQQQQQQQQQ

>DRBD2
MSENNEEQHQQQQQQQPVAVETPSAVEAPASADPSSEQSVAVEGNSEQAEDN
QGENDPSV
VPANAITGGRETSDRVLYVGNLDKAITEDILKQYFQVGGPIANIKIMIDKNNKN
VNYAFV
EYHQSHDANIALQTLNGKQIENNIVKINWAFQSQQSSSDDTKRDNNNNNNNYQQ
RRNYGNN
NRGGFRQYNSNNNNNMNMGMNMNMNMNMNNSRGMPPSSMGMP**IGAMPLP
SQGQPQQ**SQTI
GLPPQVNPQAVDHIIRSAPPRVTTAYIGNIPHFATEADLIPLFQNFGFILDFKHYPE
KGC
CFIKYDTHEQAAVCIVALANFPFQGRNLRTGWGKERSNFMPQQQQQGGQPLIM
NDQQQPV
MSEQQQQQQQQQQQ

>DPrD
MSENNEEQHQQQQQQQPVAVETPSAVEAPASADPSSEQSVAVEGNSEQAEDN
QGENDPSV
VPANAITGGRETSDRVLYVGNLDKAITEDILKQYFQVGGPIANIKIMIDKNNKN
VNYAFV
EYHQSHDANIALQTLNGKQIENNIVKINWAFQSQQSSSDDTFNLFVGDLNVNV
DDETLRN
AFKDFPSYLSGHVMWDMQTGSSRGYGFVSFTSQDDAQNAMDSMQGQDLNGR
PLRINWAAS
QTIGLPPQVNPQAVDHIIRSAPPRVTTAYIGNIPHFATEADLIPLFQNFGFILDFKH
YPE
KGCCFIKYDTHEQAAVCIVALANFPFQGRNLRTGWGKERSNFMPQQQQQGGQ
PLIMNDQQ
QPVMSEQQQQQQQQQQQ

>DPS
MSENNEEQHQQQQQQQPVAVETPSAVEAPASADPSSEQSVAVEGNSEQAEDN
QGENDPSV
VPANAITGGRETSDRVLYVGNLDKAITEDILKQYFQVGGPIANIKIMIDKNNKN
VNYAFV
EYHQSHDANIALQTLNGKQIENNIVKINWAFQSQQSSSDDT**IGAMPLPSQGQP**
**QQ**SQTIG
LPPQVNPQAVDHIIRSAPPRVTTAYIGNIPHFATEADLIPLFQNFGFILDFKHYPEK
GCC
FIKYDTHEQAAVCIVALANFPFQGRNLRTGWGKERSNFMPQQQQGGQPLIMN
DQQQPVM
SEQQQQQQQQQQQ

**Yeast growth and manipulation**

Different reagents and conditions were used to manipulate yeast strains. The following list summarize them:

Background strain growth medium: Complete Synthetic Medium
Selective medium for cloned strains: Uracil Dropout
Non-inducing conditions: glucose 2%
Inducing conditions: galactose 2%
Agar growth conditions: 30ºC
Liquid growth conditions: 30ºC, 200 rpm
Overnight: 16 hours

Number of cells was measured with a spectrophotometer reading optical density (OD) at 600 nm in all cases.

**Pre-growth (for all experiments)**

Yeast strains were thawed in agar plates from frozen stocks. After two days at 30ºC, colonies were inoculated in liquid culture. Overnight growth (30ºC, 200 rpm) in non-inducing conditions until saturation was followed by fresh dilution (~0.2 OD) in non-inducing conditions. After ~6-8 hours of growth in same conditions, cells reached late exponential phase (up to ~0.8 OD). Strains were again diluted (according to corresponding doubling times) to be ready for subsequent experimental procedures.

**Overexpression over time**

Diluted strains were grown for an overnight in either non-inducing or inducing conditions until late exponential phase. Resultant cultures (pre-induction 0 h, from overnight in non-inducing; pre-induction 16 h, from overnight in inducing) were split and used for immediate growth assessment or longer inductions.

One part of pre-induced cultures (0 and 16 h) were diluted to ~0.2 for longer induction during 8 hours (pre-induction 8 h, from previous 0 h; and pre-induction 24 h, from previous 16 h). These cultures were then used for phenotypes quantification, FRAP, growth curves assay and flow cytometry.

**Phenotypes quantification**

Strains were grown during corresponding inducing times keeping culture density always below 1 OD (diluting accordingly). Cells in exponential phase were imaged under 63× magnification on a confocal TCS SP8 microscope (Leica).

Phenotypes were classified according to condensates size in three categories: diffuse (no condensates), small (<1 µm condensates) and big (≥1 µm condensates), using ImageJ software. Quantification of phenotypes was assessed in a minimum amount of 100 cells from at least three different biological replicates for each strain and condition.

**Co-localization assays**

Strains from GFP collection transformed with Pub1-cherry p416-Gal1 plasmid were used to study localization of Pub1 full condensates. Cells were induced for 8-10 hours and imaged at exponential phase under 100× magnification on a DMRE fluorescence microscope with PRIOR Lumen 200 light (Leica).

Nucleus co-localization with ΔPS variant was also investigated. ΔPS variant fused to eGFP was induced 8 and 24 hours keeping culture density always below 1 OD (diluting accordingly). Two different nuclei staining dyes (Hoecht 1000× and DAPI 1000×) were added to liquid cultures 1 hour prior to imaging. Same fluorescent microscope and settings were used.

**Immunofluorescence assay**

Strains expressing HA-tagged proteins were induced during 8 hours until exponential phase and then fixed with 4% formaldehyde for 1 h. Cell walls were digested with ß-mercaptoethanol and zymolase (5 mg/ml). Obtained spheroplasts were permeabilized in PBS with 0.05% Tween-20 and loaded into optical-bottom 96-well plates. After blocking (BSA 1 mg/mL in PBS), fixed spheroplasts were incubated with anti-HA 3F10 rat antibody (Roche 11867431001, 1:4,000 for 1 h at room temperature), washed and incubated with secondary anti-rat Alexa Fluor 488 (Invitrogen A11006, 1:10,000 for 1 h at room temperature in the dark). Samples were finally washed in PBS, mounted in propyl gallate solution with DAPI and stored at 4°C until visualization. Cells were imaged under 100× magnification on a DMRE fluorescence microscope with PRIOR Lumen 200 light (Leica).

**Fluorescence recovery after photo-bleaching (FRAP)**

Strains were grown during corresponding inducing times keeping culture density always below 1 OD (diluting accordingly). Cells were then imaged under a Confocal TCS SP5 microscope (Leica) where bleaching was achieved with 488 Laser Power at 70% for five frames (1.3 s/frame) while recovery was recorded for 100 frames. The curves, following the fluorescence intensity of a certain cell region (i.e. cytosol and granule), were then fitted to a single exponential, following normalization with extracellular background subtraction.

**Growth assays: Toxicity and Recovery**

**Spot dilution**

Pre-induced cultures were serially diluted (six dilutions 1:5). 4 µL drop of each dilution were plated in inducing (TOXICITY) and non-inducing (RECOVERY*) agar plates. Plates were incubated for three days at 30ºC.

**Growth curves**

Pre-induced cultures were diluted to 0.1 OD in inducing conditions (TOXICITY) and non-inducing (RECOVERY). Three technical replicates per strain and condition were loaded in a 96-well plate. Growth was monitored at 30°C during two days inside the Infinite M200 microplate reader (Tecan). Note that two different set of samples were started at different times in different readers (one set includes pre-inductions 0 and 16 h, another set includes pre-inductions 8 and 24 h).

Growth curves (OD versus time) were fitted using a smoothing of second order (10 points) with Prism program. Blank OD (medium) was subtracted. Three parameters were calculated to compare strains curves:

- Lag times: measured at the point in which starts the acceleration phase, right before the exponential phase

- Doubling times (DT = LN(2)/slope): calculated adjusting a second order regression at the linear region of the exponential phase.

- Saturation OD: measured at the maximum absorbance point.

Mean and standard deviation of three biological replicates, each with three technical replicates, were computed.

* Samples from pre-induction 0 h assessed in non-inducing conditions were considered as growth control.

**Flow Cytometry**

Strains were treated as in fitness assays. ~$10^6$ cells in 1 mL PBS were used for flow cytometry. Cells were incubated with DAPI dye (1000×) to stain dead cells 5 minutes prior to analyse them.

Analysis was performed at a medium flow rate in a BD LSR II flow cytometer. The signal coming from cell doublets and aggregates was eliminated on the basis of the forward scatter area and height. Populations were separated on the basis of fluorescence intensity. 10,000 events from live cells population were collected for each sample.

### Protein extractions

100 mL culture at ~0.7 OD were pelleted and stored at -80ºC for posterior lysis. Following steps of the procedure were carried at ~0-4ºC. Protein fractions were extracted mechanically by strong vortexing (disruptor) for 10 mins with glass beads (425-600 µm) in lysis buffer (Tris HCl 1M, NaCl 5M, EDTA 0.5M, pH 7.4) and freshly added proteases inhibitors (PMSF 100×). Lysates were cleared by centrifugation (3,000 g, 10 mins) to obtain TOTAL PROTEIN extractions. Extractions were ready for western blotting, soluble/insoluble fractioning or immunoprecipitation.

Total protein extractions were used to separate soluble and insoluble fractions. Samples were transferred to polycarbonate tubes (Beckman Coulter 343776) for ultracentrifugation (55,000 rpm, 1h in a TLA 120.1 rotor). Supernatants, as SOLUBLE PROTEIN FRACTION, were mixed with 4× loading dye (Invitrogen NP0007) to proceed with western blot or store at -20ºC. Pellets, as INSOLUBLE PROTEIN FRACION, were resuspended with same concentrated loading dye and then mixed with corresponding amount of lysis buffer to proceed with western blot or store at -20ºC.

### Western blot

Protein extractions mixed with loading dye were boiled for 5 mins. Samples were run in precast NOVEX NuPAGE 4%–12% gels in denaturing conditions. The Invitrogen iBlot system was used to transfer proteins to PVDF membranes. After blocking, membranes were incubated overnight at 4°C with anti-GFP rabbit antibody (Santa Cruz sc-8334), anti-HA 3F10 rat antibody (Roche 11867431001) or anti-PGKD1 mouse antibody (Novex 459250) diluted 1:1,000, 1:4,000 and 1:10,000, respectively. Secondary incubation with anti-Protein G HRP conjugated (Millipore 18-161) at 1:10,000 was performed at room temperature during 1 h. ImageJ software was used to quantify protein bands.

### Immunoprecipitation

Total protein extractions from HA-tagged Pub1 variants were incubated with anti-HA magnetic beads (Pierce 88836; 350 µg beads per 500 µg sample) overnight at 4ºC with mixing. Beads with sample were washed three times with washing buffer (50 mM Tris/HCl (pH 7.4), 150 mM NaCl), dried at room temperature for 5 minutes and snap-frozen in liquid nitrogen.

**Mass spectrometry procedure and analysis**

Beads were re-suspended in 50 µl 8M urea, 50 mM Tris/HCl (pH8.5), reduced with 10 mM DTT for 30 min and alkylated with 40 mM chloroacetamide for 20 min at 24°C. Urea was diluted to a final concentration of 2M with 25 mM Tris/HCl (pH8.5), 10% acetonitrile and proteins were digested with trypsin/lysC mix (mass spec grade, Promega) overnight at 24°C. Acidified peptides (0.1% trifluoroacetic acid) were desalted and fractionated on combined C18/SCX stage tips (3 fractions). Peptides were dried and resolved in 1% acetonitrile, 0.1% formic acid.

Liquid chromatography coupled with tandem mass spectrometry (LC-MS/MS) was performed on a Q Exactive Plus equipped with an ultra-high pressure liquid chromatography unit (Easy-nLC1000) and a Nanospray Flex Ion-Source (all three from Thermo Fisher Scientific, Waltham, MA). Peptides were separated on an in-house packed column (100 µm inner diameter, 30 cm length, 2.4 µm Reprosil-Pur C18 resin [Dr. Maisch GmbH, Germany]) using a gradient from mobile phase A (4% acetonitrile, 0.1% formic acid) to 30% mobile phase B (80% acetonitrile, 0.1% formic acid) for 60 min followed by a second step to 60% B for 30 min, with a flow rate of 300 nl/min. MS data were recorded in data-dependent mode selecting the 10 most abundant precursor ions for HCD with a normalized collision energy of 27. The full MS scan range was set from 350 to 2000 m/z with a resolution of 70,000. Ions with a charge ≥2 were selected for MS/MS scan with a resolution of 17,500 and an isolation window of 2 m/z. The maximum ion injection time for the survey scan and the MS/MS scans was 80 ms, and the ion target values were set to 3e6 and 1e5, respectively. Dynamic exclusion of selected ions was set to 60 s. Data were acquired using Xcalibur software (Thermo Fisher Scientific).

MS raw files from 5 biological replicates of different Pub1 variants and background samples were analyzed with Max Quant (version 1.5.3.30)[16] using default parameters. Enzyme specificity was set to trypsin and lysC and a maximum of 2 missed cleavages were allowed. A minimal peptide length of 7 amino acids was required. Carbamidomethyl-cysteine was set as a fixed modification, while N-terminal acetylation and methionine oxidation were set as variable modifications. The spectra were searched against the UniProtKB yeast FASTA database (downloaded in July 2018, 6721 entries) for protein identification with a false discovery rate of 1%. Unidentified features were matched between runs in a time window of 2 min. In the case of identified peptides that were shared between two or more proteins, these were combined and reported in protein group. For label-free quantification (LFQ), the minimum ratio count was set to 1.

Bioinformatic data analysis was performed using Perseus (version 1.5.2.6)[17]. Hits in three categories (false positives, only identified by site, and known contaminants) were excluded from further analysis. Samples were grouped into Pub1 full, ΔPrLD, ΔPS and background. The data was filtered for proteins having at least 3 valid LFQ values in at least one group (pulldowns and background). Missing LFQ values were imputed on the basis of normal distribution with a width of 0.3 and a downshift of 1.5. Proteins enriched in Pub1 variants over background control were identified by two-sample t-test at different permutation-based FDR cutoffs (0.001 and 0.01) and s0 = 0.3.

**Analysis of protein interactors**

Properties of Pub1 protein interaction networks were analysed using multiCM (from Klus et al., 2015 [18]). Interactors were compared with a yeast stress granule dataset from Mitchell *et al.*, 2013 [19] and total yeast proteome (UP000002311 from UniProt [1] release 2018/11).

Other information about interactors were retrieved from corresponding on-line databases: sequence size (number of amino acids) from Uniprot ([1]; abundance from PaxDB: GPM, Aug 2014 [20]; interaction capacity BioGRID [21].

GO analysis was performed using Proteomaps. Other tools such as Gorilla and Panther were used obtaining similar results. Proteomaps was also used for visual representation.

Interactors of Pub1 variants were scanned for dosage sensitivity and essentiality using corresponding datasets. Interactors list was also compared with known Pub1 interactors, retrieved from BioGRID.

1. The UniProt C. UniProt: the universal protein knowledgebase. *Nucleic Acids Res* **45**, D158-D169 (2017).

2. Jain S, Wheeler JR, Walters RW, Agrawal A, Barsic A, Parker R. ATPase-Modulated Stress Granules Contain a Diverse Proteome and Substructure. *Cell* **164**, 487-498 (2016).

3. Alberti S, Halfmann R, King O, Kapila A, Lindquist S. A systematic survey identifies prions and illuminates sequence features of prionogenic proteins. *Cell* **137**, 146-158 (2009).

4. Pache RA, Babu MM, Aloy P. Exploiting gene deletion fitness effects in yeast to understand the modular architecture of protein complexes under different growth conditions. *BMC Syst Biol* **3**, 74 (2009).

5. Steinmetz LM*, et al.* Systematic screen for human disease genes in yeast. *Nat Genet* **31**, 400-404 (2002).

6. Ghaemmaghami S*, et al.* Global analysis of protein expression in yeast. *Nature* **425**, 737-741 (2003).

7. Sopko R*, et al.* Mapping pathways and phenotypes by systematic gene overexpression. *Mol Cell* **21**, 319-330 (2006).

8. Makanae K, Kintaka R, Makino T, Kitano H, Moriya H. Identification of dosage-sensitive genes in Saccharomyces cerevisiae using the genetic tug-of-war method. *Genome Res* **23**, 300-311 (2013).

9. Hentze MW, Castello A, Schwarzl T, Preiss T. A brave new world of RNA-binding proteins. *Nat Rev Mol Cell Biol* **19**, 327-341 (2018).

10. El-Gebali S*, et al.* The Pfam protein families database in 2019. *Nucleic Acids Res* **47**, D427-D432 (2019).

11. Potter SC, Luciani A, Eddy SR, Park Y, Lopez R, Finn RD. HMMER web server: 2018 update. *Nucleic Acids Res* **46**, W200-W204 (2018).

12. Bolognesi B*, et al.* A Concentration-Dependent Liquid Phase Separation Can Cause Toxicity upon Increased Protein Expression. *Cell reports* **16**, 222-231 (2016).

13. Lancaster AK, Nutter-Upham A, Lindquist S, King OD. PLAAC: a web and command-line application to identify proteins with prion-like amino acid composition. *Bioinformatics* **30**, 2501-2502 (2014).

14. Janke C*, et al.* A versatile toolbox for PCR-based tagging of yeast genes: new fluorescent proteins, more markers and promoter substitution cassettes. *Yeast* **21**, 947-962 (2004).

15. https://yeastgfp.yeastgenome.org/.

16. Cox J, Mann M. MaxQuant enables high peptide identification rates, individualized p.p.b.-range mass accuracies and proteome-wide protein quantification. *Nat Biotechnol* **26**, 1367-1372 (2008).

17. Tyanova S*, et al.* The Perseus computational platform for comprehensive analysis of (prote)omics data. *Nat Methods* **13**, 731-740 (2016).

18. Klus P, Ponti RD, Livi CM, Tartaglia GG. Protein aggregation, structural disorder and RNA-binding ability: a new approach for physico-chemical and gene ontology classification of multiple datasets. *BMC Genomics* **16**, 1071 (2015).

19. Vance C*, et al.* ALS mutant FUS disrupts nuclear localization and sequesters wild-type FUS within cytoplasmic stress granules. *Hum Mol Genet* **22**, 2676-2688 (2013).

20. https://pax-db.org/.

21. https://thebiogrid.org/.
